# Fluorescence Anisotropy Analysis of the Interaction between Doxorubicin and DNA Origami Nanostructures

**DOI:** 10.1101/2024.06.20.599777

**Authors:** Ekaterina S. Lisitsyna, Anna Klose, Elina Vuorimaa-Laukkanen, Heini Ijäs, Tatu Lajunen, Klaus Suhling, Veikko Linko, Timo Laaksonen

## Abstract

Owing to doxorubicin’s high DNA binding affinity, doxorubicin-loaded DNA origami nanostructures (DOX-DONs) are promising nanocarriers against cancer. However, understanding the interactions between doxorubicin (DOX) and DNA origami nanostructures (DONs) is important to ensure the quality of DOX-DONs. This interaction is often taken for granted and the influence of DOX loading conditions is poorly characterized. Exploiting the inherent fluorescence of DOX, steady-state and time-resolved fluorescence anisotropy spectroscopy techniques are used for characterizing non-destructively the binding between DOX and DONs, and the purity of formed complexes. The difference in fluorescence anisotropy between free DOX and DOX-DONs confirms the DOX-DON complex formation. Further, at loading ratios of DOX to DNA base pairs > 0.5, homo-Förster resonance energy transfer (homo-FRET) between closely packed DOX molecules is observed. Moreover, time-resolved anisotropy reveals DOX aggregation on DONs at high loading ratios > 1. For loading ratios > 0.1, spin-filtration to remove excess free DOX is efficient and necessary, though at loading ratios > 1 some DOX aggregates remain attached to the DONs. In summary, fluorescence anisotropy analysis provides more detailed information and insight into DOX-DONs compared to regularly used fluorescence intensity-based characterization methods, and these results can help designing more efficient and safer DNA intercalator-based nanocarriers.

## 1. Introduction

DNA nanotechnology is based on harnessing the sequence-specific hybridization of DNA molecules into DNA nanostructures through self-assembly.^[1]^ DNA origami nanostructures (DONs), in particular, form through cooperative binding of many short, custom-designed DNA staple strands to base-complementary sections of a single-stranded scaffold strand, that guides the assembly into 2D or 3D DONs.^[2–4]^ Their properties can be precisely tuned and customized,^[5–7]^ making DONs attractive for various applications, including drug delivery.^[6,8]^ While the structural integrity of DONs under physiological conditions can be challenging to maintain due to their susceptibility to enzymes and low-cation concentrations,^[9,10]^ coating and design strategies may improve their stability.^[11,12]^ Especially their biodegradability and biocompatibility makes them attractive nanostructures in biomedical applications.^[13,14]^

As nanocarriers, DONs can host various therapeutic molecules, such as antibody fragments,^[15]^ nucleic acids,^[16–18]^ enzymes,^[19]^ or small molecules such as doxorubicin.^[20–22]^ The chemotherapeutic agent doxorubicin (DOX, SI0) is a fluorescent DNA-intercalator, commonly used to assess DONs as drug carriers. DOX damages DNA in cells through the inhibition of the DNA topoisomerase II enzyme,^[23]^ and dysregulating other processes.^[24]^ This, however, also causes clinical side effects like cardiotoxicity, requiring new strategies to reduce off-target DOX exposure and toxicity.^[25]^ Against this backdrop, DOX-loaded DONs (DOX-DONs) showed promising in vitro and in vivo results, overcoming drug resistance via targeting strategies and combination therapy,^[21,26,27]^ and inhibiting tumor growth in mice with minimal toxicity,^[18,22,28]^ making them attractive nanocarriers for further characterizations.

DOX intercalates DNA by inserting itself planarly between DNA bases. At higher concentrations, DOX starts aggregating on top of DNA,^[29,30]^ or DOX aggregates precipitate due to the high pH and ionic strength of buffers commonly used for DON loading.^[31–33]^ Yet, reported loading conditions are little-characterized, and vary regarding DOX concentration, temperature, buffers, incubation times and purification steps, leading to overestimations of DOX loaded into DONs^[31]^: This is prominent for DOX-DONs that are recovered after loading through centrifugation, because DOX aggregates precipitate alongside DOX-DONs and only soluble DOX is removed with the supernatant.^[21,22,26,27,34,35]^ Some protocols avoid this by using spin-filtration.^[18,20,28,36]^ In consequence, the amount of DOX loaded, usually indirectly quantified in the supernatant,^[21,22,26,35]^ or the filtrate,^[28]^ will be overestimated if the effectiveness of purification is not established, and further efficacy studies will be based on false premises. Hence, it is important to choose and verify appropriate loading and purification conditions for successful DOX-loading into DONs.

Many commonly used methods for quantifying the composition of the DOX-DONs have likewise limitations. Through absorption measurement, all DOX in purified DOX-DON samples is detectable,^[27,34,36]^ but because of the spectral overlap between free DOX and DNA-bound DOX, this cannot shed light on the state of DOX. Similarly, quenching of DOX-fluorescence upon DNA binding,^[21]^ reduced migration of DOX-DONs in gel electrophoresis-based mobility shift assays,^[20,27,36]^ and visual confirmation through imaging^[20]^ only provide qualitative confirmation of binding and structural integrity of the DOX-DONs. Destruction of DOX-DONs by heating or enzymatic digestion to release and quantify the DOX remains similarly biased to the presence of unbound DOX.^[18,31,36]^ Hence, more advanced spectroscopic techniques are needed to elucidate the state of DOX in DOX-DON samples.

Fluorescence techniques are non-destructive and highly sensitive, making them ideal for characterizing intricate biological samples and nanoparticles,^[37,38]^ whilst preserving their structural and functional integrity. A set of parameters, including intensity, wavelength, lifetime, and polarization characterize fluorescence, all of which can vary sharply through interactions of the fluorophore with the local environment.^[39]^ For instance, fluorescence lifetime imaging (FLIM) has been successfully used to monitor the cellular uptake of DOX,^[40]^ and its induction of apoptosis.^[41]^

Furthermore, in fluorescence anisotropy measurements, exciting a fluorophore with polarized light results in similarly polarized emission, which, however, is reduced if the fluorophore can freely rotate in solution or depolarize by non-radiative energy transfer. The fluorescence polarization is expressed by the anisotropy value (*r*), and provides information otherwise unobtainable from emission spectra, intensity or lifetime measurements: these include molecular orientation, aggregation, rotational diffusion and energy migration among chemically identical molecules (homo-FRET, homo-Förster resonance energy transfer).^[42–46]^

Fluorescence anisotropy has been used to probe nanoparticle size,^[47–49]^ binding and conformational dynamics of biomolecules, such as proteins, nucleic acids, and aptamers,^[45,49– 52]^ as well as chromophore organization and energy transfer.^[53–56]^ In respect to the latter, fluorescence anisotropy is the only available method to detect homo-FRET.^[57,58]^ Homo-FRET can be used for ion-sensing^[54]^ and transferring energy over long distance via photonic wires,^[59,60]^ but moreover, it can reveal protein dimerization,^[56]^ cluster sizes,^[61–63]^ and the distance,^[40]^ as well as loading and packing density of fluorophores.^[41]^ Exploiting the intrinsic fluorescence of DOX, fluorescence lifetime and anisotropy measurements can be used to characterize DOX and its interactions. Fluorescence anisotropy has been used for characterizing DOX itself,^[64]^ its localization and incorporation into different formulations,^[65–69]^ and its interaction, binding and release behaviors from other macromolecules.^[70–72]^ In this work, we demonstrate the applicability of steady-state and especially time-resolved fluorescence anisotropy spectroscopy for studying DONs loaded with DOX.

## 2. Results and Discussion

The interactions between DOX and DONs were studied both for unpurified and purified samples. For the latter, after loading DOX into DONs, spin-filtration was used to remove the excess DOX (**Figure 1**). With steady-state and time-resolved fluorescence anisotropy spectroscopy we studied eight different loading ratios [DOX]/[bp_DNA_] (**Table 1**) to reveal how the amount of DOX influences the DOX-DON complex formation and the success of the purification.

**Table 1.**
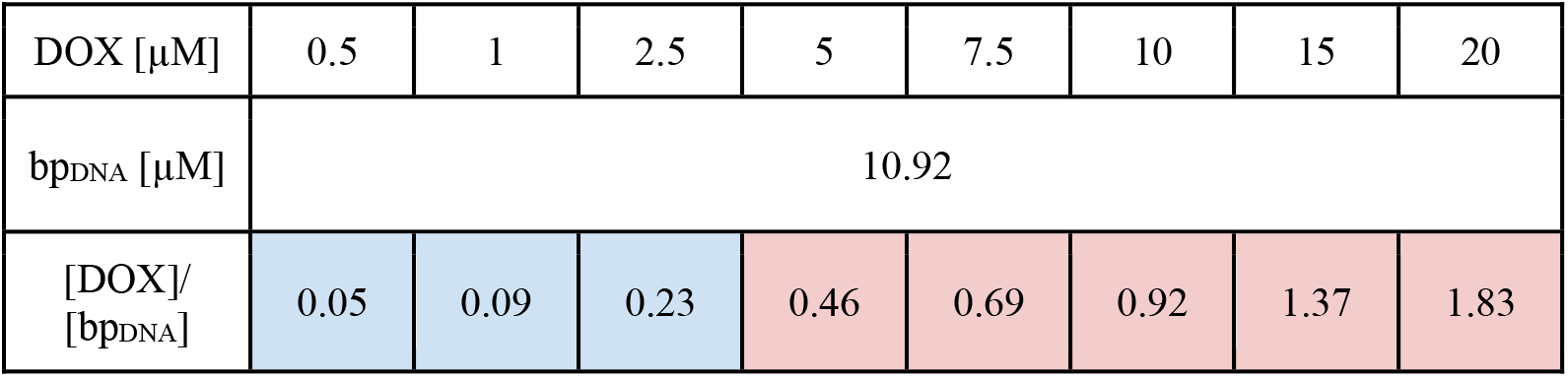
Loading ratio [DOX]/[bp_DNA_] between the concentration of DOX (µM) and the concentration of DNA base pairs in DONs (bp_DNA_; µM) in the loading reaction before purification. Low loading ratios < 0.3 are highlighted in blue, higher loading ratios in red.

| DOX [$\mu\text{M}$] | 0.5 | 1 | 2.5 | 5 | 7.5 | 10 | 15 | 20 |
| --- | --- | --- | --- | --- | --- | --- | --- | --- |
| $\text{bp}_{\text{DNA}}$ [$\mu\text{M}$] | 10.92 | | | | | | | |
| $[\text{DOX}]/[\text{bp}_{\text{DNA}}]$ | 0.05 | 0.09 | 0.23 | 0.46 | 0.69 | 0.92 | 1.37 | 1.83 |

**Figure 1.**
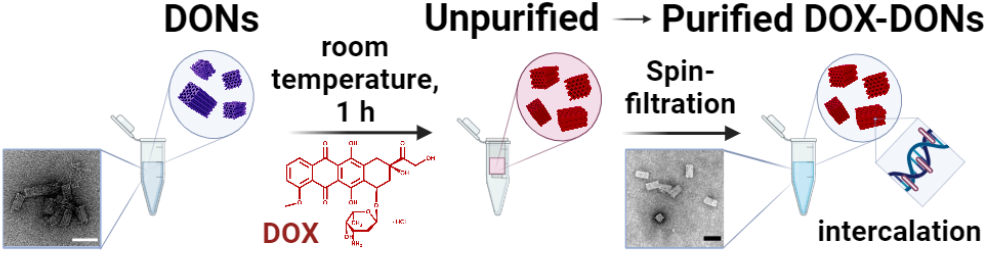
Loading of DONs with DOX in water and further purification. Scale bars in TEM images are 50 nm.

### 2.1. Emission and Excitation Spectra

For free DOX in deionized water, the excitation spectrum had two maxima at 476 and 495 nm, the first one being slightly more intensive (**Figure SI6**). Upon binding to the DONs, the fluorescence intensity of DOX decreased because of quenching,^[21,73,74]^ and the excitation spectra showed a clear shape change that depended on the loading ratio: For unpurified DOX-DONs the excitation spectra resembled that of free DOX with two maxima at 478 and 496 nm at loading ratios > 0.3, but for purified DOX-DONs only at loading ratios > 1. However, for purified DOX-DONs at loading ratios < 1, excitation spectra had one maximum at 499 nm and two shoulders at around 480 nm and 550 nm, indicating that DOX binds to the DONs and excess DOX was efficiently removed (**Figure 2a**). The little difference in excitation spectra between DOX-DONs before and after purification at loading ratios < 0.1, suggested complete DOX-loading into the DONs or only a little free DOX left in the unpurified samples (**Figure SI6, e**).

**Figure 2.**
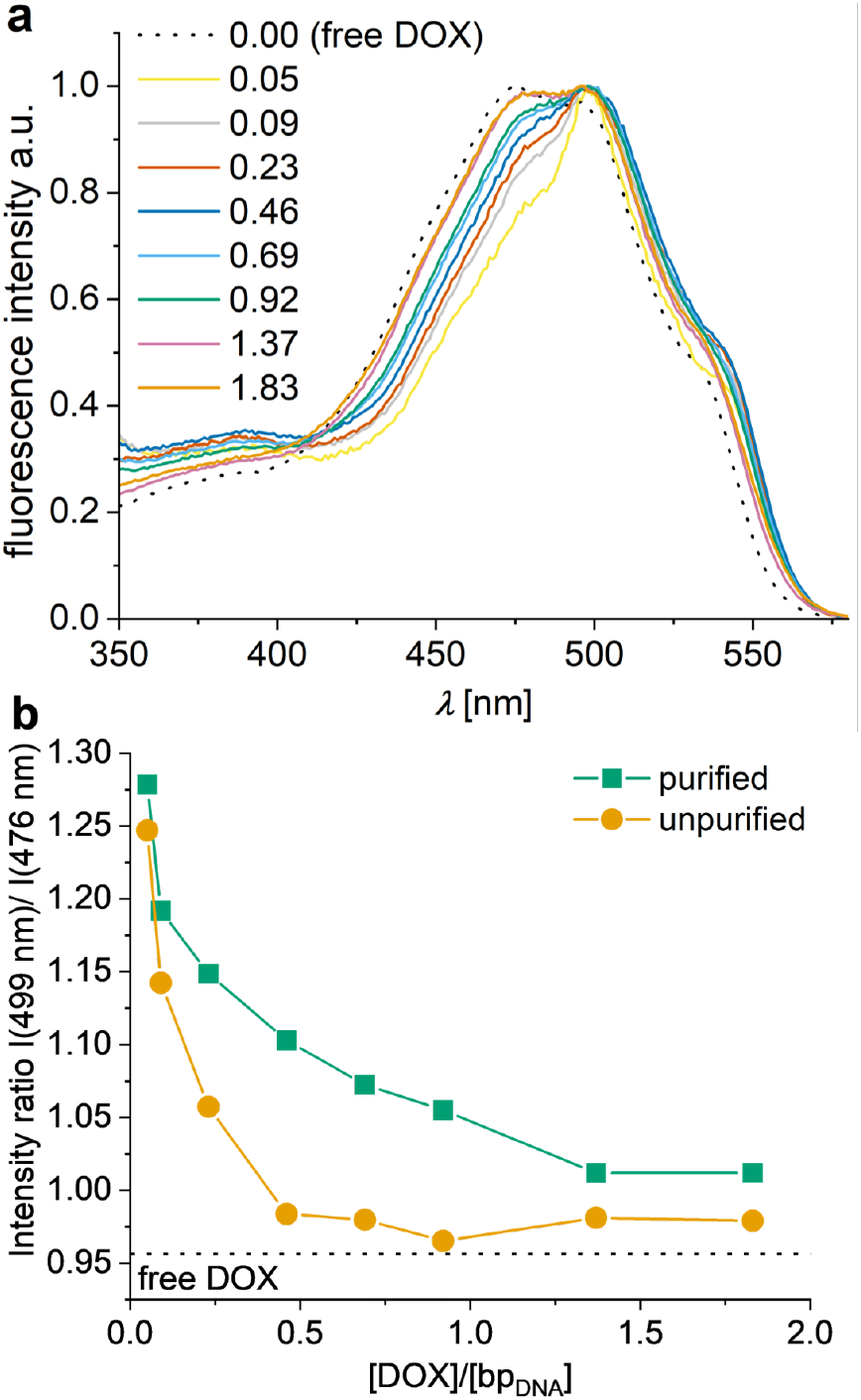
a – Normalized excitation spectra of purified DOX-DONs at different [DOX]/[bp_DNA_] loading ratios. b – Ratio of the fluorescence intensities *I*(499 nm)/*I*(476 nm) taken from the excitation spectra of free DOX, purified and unpurified DOX-DONs at different [DOX]/[bp_DNA_] loading ratios. *λ*_exc_ = 483 nm, *λ*_det_ = 600 nm. (ratio for free DOX (5 µM) shown as a dotted line).

To illustrate these changes in the excitation spectra as a result of DOX-DON complex formation and purification, the ratio of intensities at the main peak of DNA-bound DOX (499 nm) and of free DOX (476 nm) was plotted as a function of loading ratio (**Figure 2b**). The almost similar intensity ratios for loading ratios < 0.1 demonstrated that DONs complexed almost all DOX molecules, and purification was unnecessary. For loading ratios of 0.2–1.0, the intensity ratios for purified and unpurified DOX-DONs substantially differed, confirming that the purification removes free DOX. For loading ratios > 1, however, the trends resembled each other, indicating that some free DOX could remain in the system after spin-filtration. Alternatively, DOX dimerization/aggregation at high concentrations could lead to the excitation spectrum widening.

Only small differences were observed between the emission spectra for free DOX, purified, and unpurified DOX-DONs: Free DOX had two shoulders around 550 nm and 650 nm and a maximum at 596 nm, that shifted upon DON-binding to 601 nm for both purified and unpurified DOX-DONs (**Figure SI6d, f**). Moreover, the widening of the spectra towards longer wavelengths, observed for purified DOX-DONs at loading ratios > 1, can indicate that dimerization or aggregation of DOX takes place. Thus, steady-state emission and excitation spectra can be used to monitor the loading and purification of DOX-DONs, although for loading ratios > 0.3 it still remains unclear whether the removal of free DOX through purification was sufficient. As the excitation and emission spectra alone do not provide clear and full understanding of the complex formation between DOX and DON at high loading ratios, and as multiple explanations of the observed peaks are possible (described earlier), there is a need for other methodology to clarify the process.

### 2.2. Steady-State Fluorescence Anisotropy

Steady-state fluorescence anisotropy values for DOX-DONs elucidated the presence of free DOX in the samples (**Figure 3**): The anisotropy of free DOX was low, 0.04 (striped bar), but the anisotropy increased to 0.16-0.18 in purified DOX-DONs at loading ratios < 0.3 because the binding to DONs reduced the free rotation and emission depolarization of DOX. For loading ratios < 0.1, purification appeared unnecessary since the fluorescence anisotropy for the unpurified DOX-DONs (yellow bars) was similar to the purified ones (green bars). In contrast, for unpurified DOX-DONs at loading ratios > 0.1, the anisotropy decreased strongly due to the presence of free DOX in the samples. After purification, the anisotropy of these samples increased, but not to the maximum value of 0.185.

**Figure 3.**
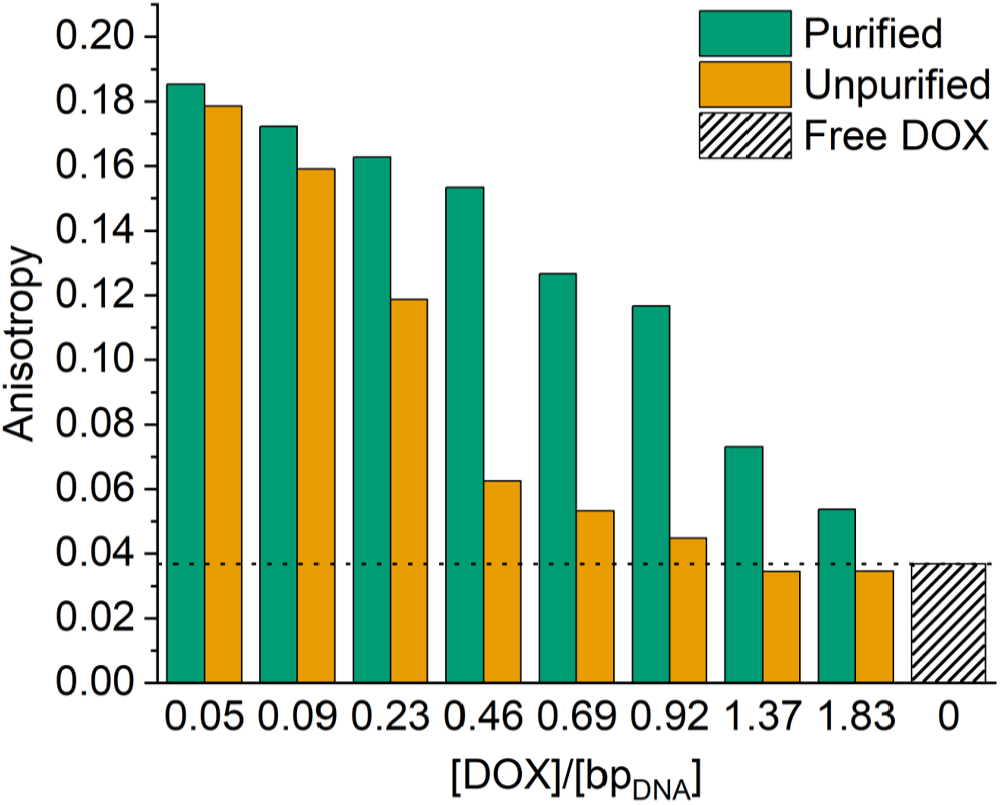
Fluorescence anisotropy of free DOX in water (5 µM, striped bar) and DOX-DONs at different loading ratios [DOX]/[bp_DNA_], for both purified (green) and unpurified (yellow) samples.

The lowered steady-state anisotropy for purified DOX-DONs at loading ratios > 0.1 may stem either from a contribution of free DOX after insufficient purification, or from nonradiative fluorescence energy transfer between DOX molecules, intercalated closely to each other in DONs. In the latter case, the fluorescence lifetime would remain the same (see Methods Section, **Equation 4**), but the energy transfer between two closely located DOX molecules (homo-FRET) would provide an additional pathway for depolarization after excitation, resulting in lower anisotropy values. While steady-state fluorescence anisotropy confirms the binding of DOX to DONs, and the successful DOX removal for loading ratios < 0.5, at higher loading ratios the efficiency of spin-filtration and the role of homo-FRET remained uncertain.

### 2.3. Fluorescence Decays

Fluorescence decay measurements clarified the low fluorescence anisotropy in purified DOX-DONs at loading ratios > 0.5 (**Figure 4a, SI7**). Free DOX in water had a mono-exponential fluorescence decay with a lifetime of 1.05 ns, in agreement with previous work (Figure 4a, black; SI7).^[64,69,75–77]^ For all purified DOX-DONs, except the one with the highest loading ratio, the decays were bi-exponential having almost equally contributing lifetimes of about 0.14 and 1.00 ns (Methods, **Equation 5**) (**Figure SI9**, green squares and bars). Likewise, the amplitude-weighted average lifetime was almost constant and significantly shorter than for free DOX (**Figure 4b**). This indicates that all purified DOX-DONs were free of excess DOX, which contradicts the previous assumption^[31]^ that after purification the DOX-DONs immediately reestablish an equilibrium with free DOX. Instead, the binding between DOX and DONs seems to be very strong. The decay for the highest loading ratio of 1.83 however deviates from the rest of purified DOX-DONs which is in agreement with the broader fluorescence spectrum observed for this sample (**Figure SI6d**) suggesting the formation of DOX dimers/aggregates on the DON matrix.

**Figure 4.**
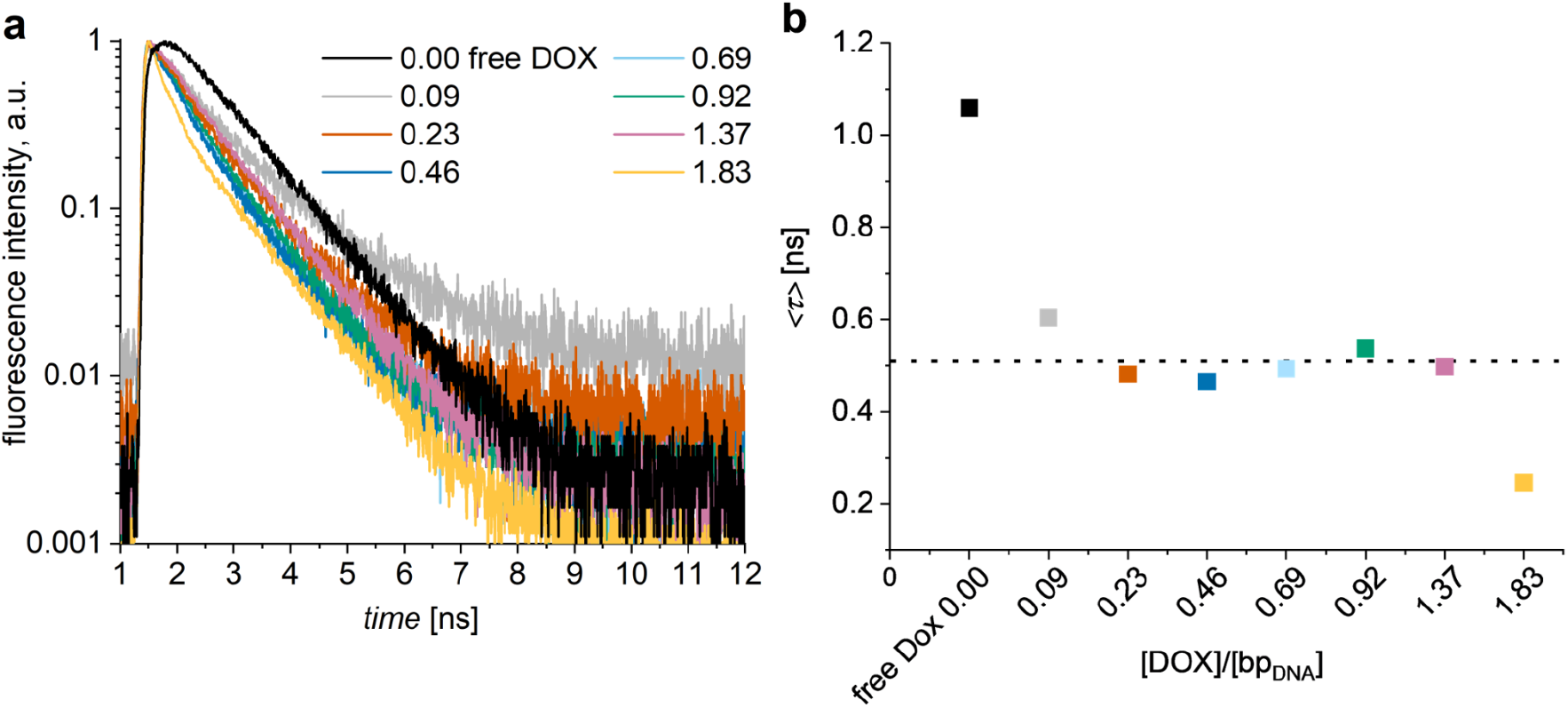
a – Normalized fluorescence decays of purified DOX-DONs at different [DOX]/[bp_DNA_] loading ratios. b – Amplitude-weighted fluorescence lifetimes calculated based on biexponential fitting of the fluorescence decays (a). *λ*_exc_ = 483 nm, *λ*_det_ = 600 nm. Average of τ for [DOX]/[bp_DNA_] from 0.09 to 1.37 is equal to 0.51 ns and shown as a horizontal dashed line.

For unpurified DOX-DONs, the fluorescence decays depend on the loading ratio: For loading ratios < 0.3, the decays were identical for both purified and unpurified DOX-DONs, but different from free DOX (**Figure 5a, SI7a, b**), indicating that all DOX molecules were bound to DONs. At loading ratios of 0.3 - 1.0, the fluorescence decays for unpurified DOX-DONs (**Figure 5b, SI7 c-e**, yellow) resembled more that of free DOX (black) than the purified DOX-DONs (green), confirming the presence of free DOX together with DOX-loaded DONs.

**Figure 5.**
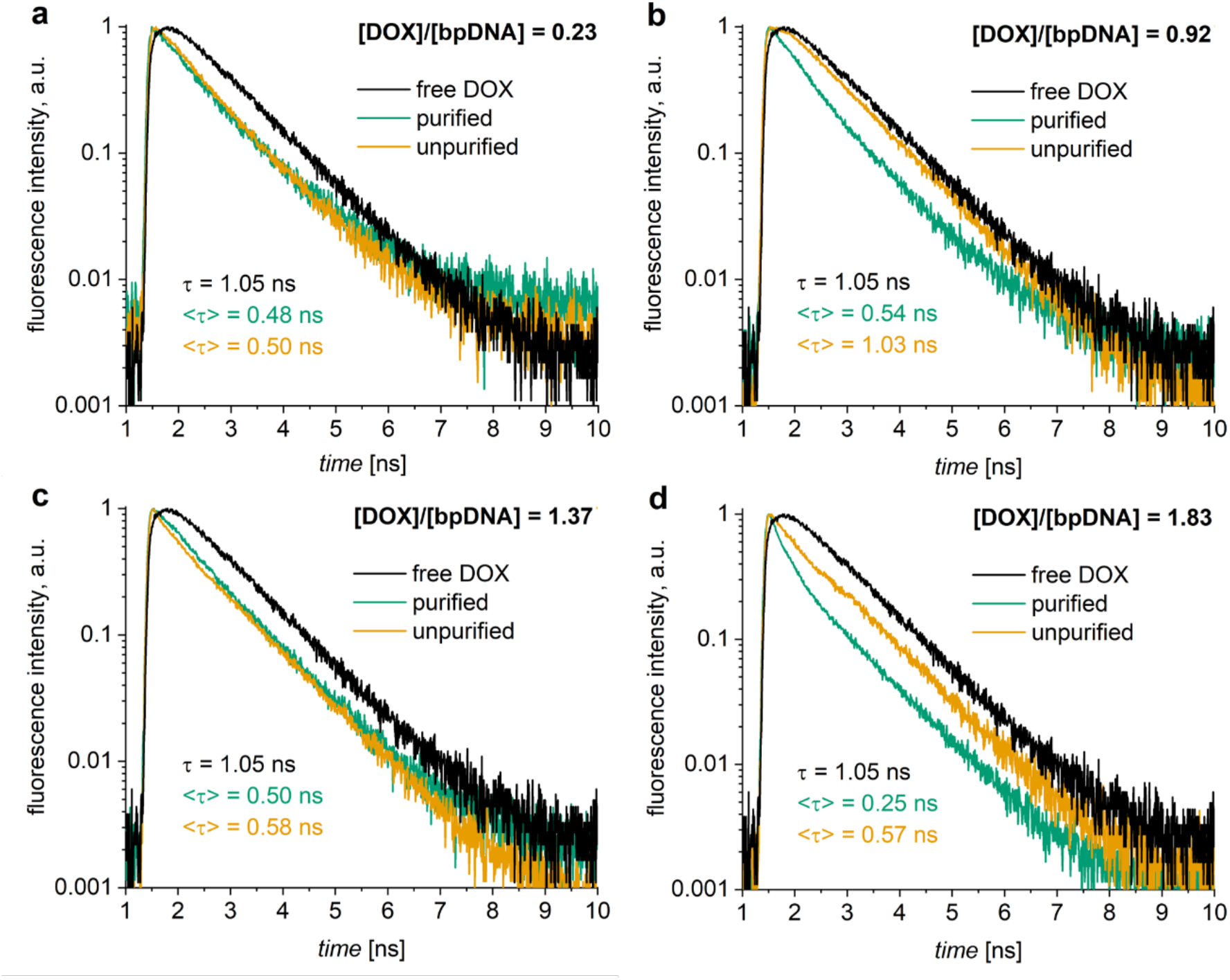
Comparison of normalized fluorescence decays of purified (green) and unpurified (yellow) DOX-DONs at different [DOX]/[bp_DNA_] ratios with that of free DOX (5 µM, black) in water, including their respective amplitude-weighted fluorescence lifetimes τ.

The situation was different for loading ratios > 1, where the unpurified DOX-DONs had an average lifetime lower than that of free DOX, suggesting the absence or reduced amount of free monomeric DOX in solution (**Figure 5c, d, SI7f, g**). Instead, fitting revealed the presence of a short-living DOX species, with a lifetime even shorter than that for DOX-DONs at loading ratios < 1 (**Table SI10**). This could indicate the presence of DOX dimers or aggregates attached to the DNA origami matrix in the samples. Free DOX dimers in water have a very short lifetime (2 ps),^[64]^ and they are invisible for the present measuring system with a time resolution of 130 ps (**Figure SI11**) but may become visible when stabilized by the DONs. Pérez-Arnaiz *et al*. previously described intercalation as the first type of complexation mechanism between DOX and ct-DNA at [DOX]/[bp_DNA_] ratios < 0.35, but at the higher ratios, DOX can aggregate on top of another DNA-intercalated DOX molecule, thus forming another type of complex.^[29]^ Another explanation of the lifetime shortening could be the energy transfer from monomeric DOX to its dimers or aggregates (hetero-FRET). The overlap between the DOX dimer absorption and its monomer fluorescence spectrum also speaks to the explanation.^[64]^ The effect is even more pronounced in the purified DOX-DON sample at the highest loading ratio of 1.83 (**Figure 5d**). It seems that the purification step was efficient in removing free DOX for the loading ratio of 1.83 as the contribution of a longer lifetime (∼ 1 ns) was smaller after purification of DOX-DONs (**Figure 5d**). DOX dimers/aggregates seemed to stay attached to the DON matrix even after spin-filtration leading to domination of a short-living lifetime component in the purified DOX-DON sample due to FRET at the highest loading ratio of 1.83 compared to the unpurified sample at the same ratio (**Table SI10**). Although, in case of [DOX]/[bp_DNA_] ratio of 1.37 (**Figure 5c**), the unpurified sample curve almost coincides with the curve for the purified sample that contrasts with all the other ratios of the studied series. Based on the above discussion we claim that free DOX in solution started dimerizing and then further aggregating on the DON matrix at the ratio of 1.37. At this ratio, the amount of DOX reached the concentration at which equilibrium shifted from the mixture of monomeric DOX in solution and DON complexes with monomeric DOX to dimeric and then aggregated DOX attached onto the DON matrix. Removal of DOX monomer from the solution by purification should lead to a shorter fluorescence lifetime. However almost unchanged lifetime after spin-filtration (**Figure 5c**) can be explained by removal of DOX from dimers on the DON matrix resulting in the fluorescence lifetime increase counterweighing the decrease due to free DOX removal. Another explanation could be that negligible amount of free DOX is left in the solution upon reaching the concentration at which DOX prefers to be in dimeric form attached to DNA and there are enough binding sites for all the added DOX to be bound. Apparently, fluorescence lifetime measurements exclusively are not enough to conclude about the purification effect on the DOX dimers/aggregates attached to DON matrix, and the system will be additionally studied by time-resolved anisotropy.

### 2.4. Confirmation of Homo-FRET

Fluorescence lifetimes ruled out the presence of free DOX in purified DOX-DONs (Figure 4), indicating that at loading ratios > 0.5 the anisotropy remained lower (**Figure 3**, green bars) just because of the homo-FRET between DOX molecules. Due to the overlap between their absorption and emission spectra (**Figure 6a**), DOX molecules can transfer excited state energy amongst each other, given they are in sufficient proximity. Taking into account the spectral overlap integral *J*(*λ*) of 2.977 × 10^13^ M^−1^ cm^−1^ nm^4^ (Methods, **Equation 2**), the orientation factor of *κ*^2^ = ⅔,^[57]^ a quantum yield of *Φ* = 0.044,^[69]^ and a refractive index *n* = 1.333 for water, the Förster radius for DOX-DOX homo-FRET is *R*_0_ = 1.7 nm (**SI12**). Thus, homo-FRET can take place with 50% efficiency if the DOX molecules are intercalated at a distance of 5 base pairs, provided the distance between the base pairs equals 0.34 nm. This is in agreement with previous reports showing that DONs can host one DOX molecule up to every 2-3 base pairs.^[31]^

**Figure 6.**
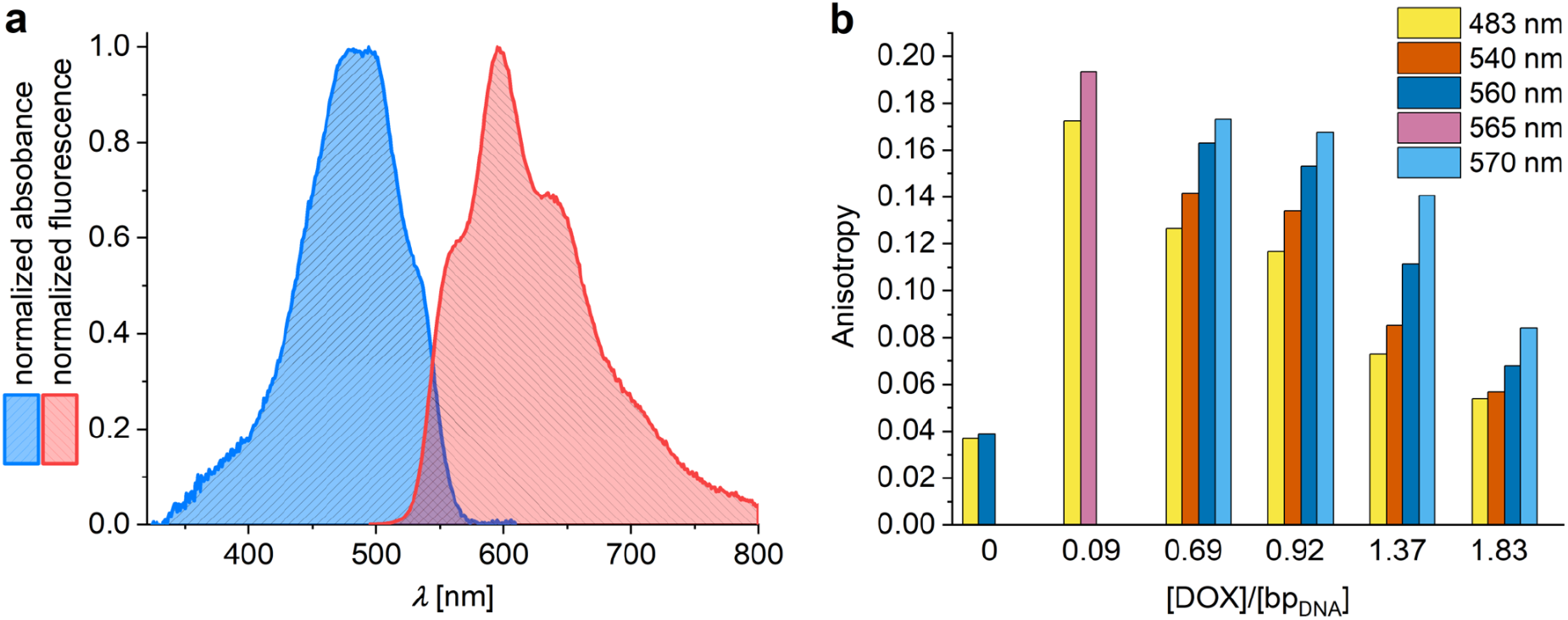
a– Absorption and emission spectra of free DOX in water showing their overlap. b – Steady-state fluorescence anisotropy measurements at different excitation wavelengths for purified DOX-DONs at [DOX]/[bp_DNA_] loading ratios of 0.09 and 0.69-1.83 in comparison with free DOX in water (5 µM).

Confirming homo-FRET with additional steady-state anisotropy measurements required excitation of purified DOX-DONs with light of the lowest possible energy, namely light at the red edge of DOX absorption, to suppress homo-FRET and prompt an increase in fluorescence anisotropy.^[56–58]^ The anisotropy of purified DOX-DONs at loading ratios 0.69 and 0.92 rose with the shift to longer excitation wavelengths (∼560–570 nm), thus, confirming the presence of homo-FRET (**Figure 6b**). The maximum fluorescence anisotropy values were almost equal to those of the DOX-DONs at lower loading ratios (< 0.3, **Figure 2**). While even for unpurified DOX-DONs some increase in anisotropy with red edge excitation was observable, the effect was less pronounced and overshadowed with increasing amounts of free DOX (**Figure SI13**).

Interestingly, upon red edge excitation of purified DOX-DONs with loading ratios > 1, the fluorescence anisotropy signal still fell short of completely recovering to its maximum value (**Figure 6b**), possibly indicating the contribution of a second type of DOX aggregate to the anisotropy value. To confirm and validate these conclusions, we performed time-resolved fluorescence anisotropy measurements on the purified and unpurified DOX-DONs.

### 2.5. Time-Resolved Fluorescence Anisotropy

Fluorescence anisotropy decays were obtained from parallel and perpendicular intensity decays (Methods, **Equation 7, Figure SI14**). For free DOX, the time-resolved anisotropy decayed to zero, indicating the free rotation of DOX (**Figure 7a**). Similarly, fitting using a monoexponential model (Methods, **Equation 8**) yielded *r*_*∞*_ ∼ 0 (free rotation) and a rotational correlation time (*θ*) of about 0.30 ns, independent of concentration within the studied range (**Figure SI15**), which is in line with previous results (**Figure 7b**, black dot).^[69]^ Using Stokes-Einstein relation (Methods, **Equation 9**), the viscosity *η* of water 1 cP, the temperature *T* of 20 °C, and the Boltzmann constant of 1.38 × 10^−23^ J K^-1^, the molecular volume of DOX *V*_*DOX*_ was found to be 1.21 nm^3^ (assuming it to be spherical) and the molecular diameter of DOX 1.32 nm, respectively. It agrees with the literature value of about 1.5 nm,^[78]^ validating the anisotropy measurement and confirming that DOX is in its monomeric form in water at the studied concentrations.

**Figure 7.**
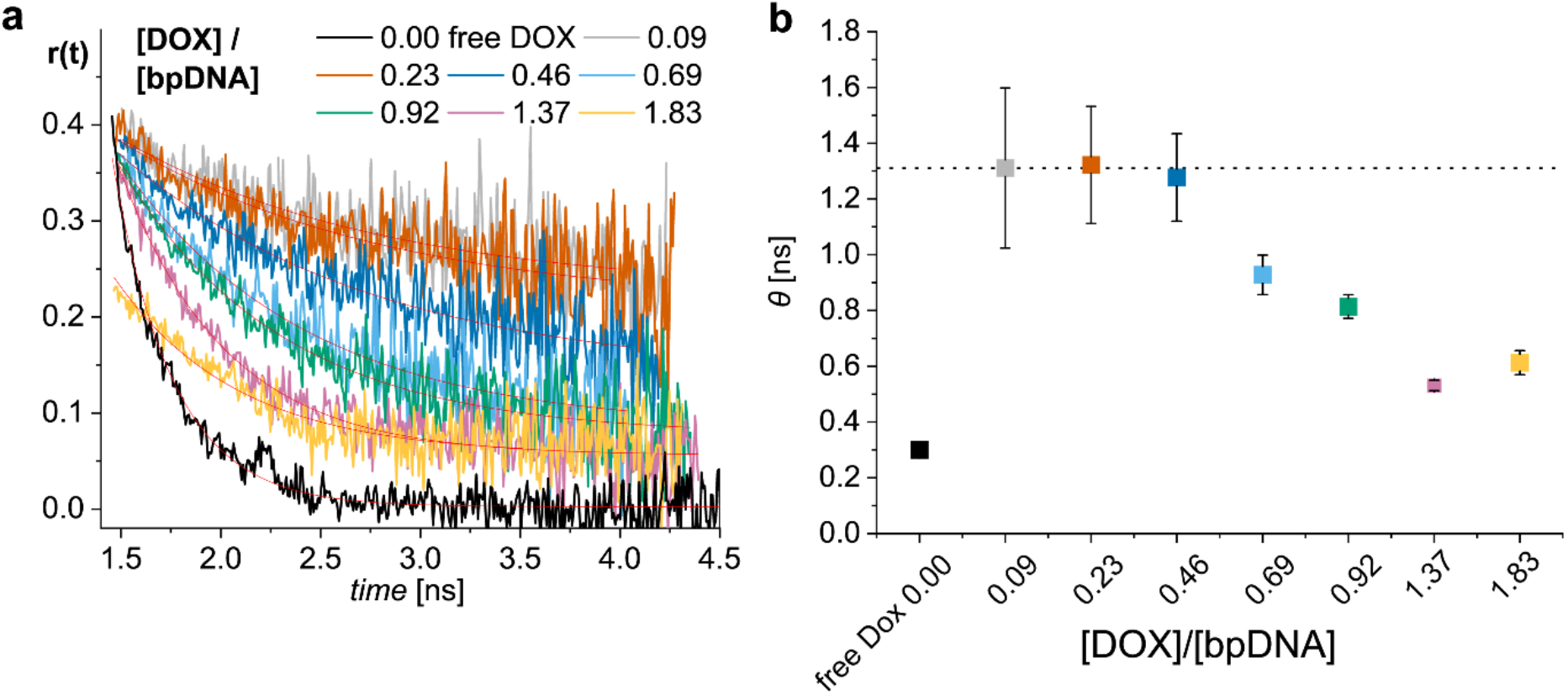
a – Time-resolved fluorescence anisotropy decays of purified DOX-DONs at different [DOX]/[bp_DNA_] loading ratios excited at 483 nm and detected at 600 nm with monoexponential fitting curves (Methods, Equation 8). b – Rotational correlation times (*θ)* calculated via the monoexponential fitting of the decays (a).

On the other hand, anisotropy decays for purified DOX-DONs leveled on different plateaus, depending inversely on the loading ratio (Figure 7a). At low loading ratios < 0.23, the rotational correlation time was much higher, as apparent in the curves shown in **Figure 7a** and in the fitted *θ* values in **Figure 7b** (**Table SI16**). Also, the non-zero positive anisotropy value of *r*_*∞*_ suggested restricted movement of DOX due to its complexation with DONs. However, for the loading ratios < 0.23, the rotational correlation times were significantly longer than the fluorescence lifetime (**Figure 4b**), reducing the accuracy of the anisotropy decay fitting. Despite the rather high experimental uncertainty of DOX rotational correlation times for DOX-DONs at low loading ratios (**Figure 7b**, dots with large error bars, **Table SI16**), it was still possible to conclude from the similarity of the anisotropy decays a common type of binding for DOX-DONs at low loading ratios (**Figure 7a**, grey, orange). Upon increasing the loading ratio to 0.46 - 1.37, the rotational correlation times decreased progressively, and the anisotropy decayed faster to lower plateaus of non-zero positive anisotropy values *r*_*∞*_ (**Figure 7a**, blue, green, pink). Similar to steady-state anisotropy (**Figure 3**), this was due to homo-FRET enabled by the dense DOX-packing into the DONs.

The short fluorescence anisotropy decays of unpurified DOX-DONs due to free DOX emphasized again the need for purification, starting from loading ratios *⩾* 0.46 (**Figure SI17**). Only the loading ratio of 0.09 displayed similar anisotropy decays for both purified and unpurified DOX-DONs. In contrast, the loading ratio of 0.23 showed similar lifetimes, but a difference in anisotropy decays before and after purification. Hence, the time-resolved fluorescence anisotropy appears to be more sensitive, allowing to detect even minute amounts of unbound DOX.

Anisotropy decays of loading ratios > 1, started from a limiting anisotropy value (*r*_0_) of about 0.22 in contrast to ∼0.39 - 0.4 characteristic for monomeric bound DOX (**Figure SI17f, g**). This unusual shape of the decays with the reduced *r*_0_ was observed for the same samples for which the unexpectedly short fluorescence intensity decays were observed (**Figure 5c, d**). This indirectly confirmed again the dimerization of DOX on the DONs, as the dimer component usually decays faster than the instrument response function and expectedly leads to a drop in the limiting anisotropy. The amplitude of the limiting anisotropy decrease is inversely proportional to the fraction of dimers in the population.^[79]^ Purification of the sample with loading ratio 1.37 recovered the *r*_0_ value back to 0.38, suggesting successful removal of DOX from dimers/aggregates attached to the DON surface. However, this was not the case for the highest loading ratio of 1.83, where the anisotropy decay remained unchanged after the purification showing that the DOX dimers/aggregates stay on DON matrix even after the spin-filtration. The result reveals the limitations of the purification efficiency at such high loading ratios and clarifies the lifetime measurements. The observations are also in line with the results of steady-state anisotropy for the loading ratios > 1. Only partial recovery of the steady-state anisotropy upon red-edge excitation for the highly loaded DOX-DONs (**Figure 6b**) can be explained by the presence of DOX aggregates allowing for a hetero-FRET between DOX in monomeric and aggregated forms. In conclusion, fluorescence anisotropy decays appeared to be more sensitive than fluorescence lifetime measurements in confirming DOX-DON binding and free DOX, especially elucidating that DOX dimers at higher loading ratios (> 1) allow for hetero-FRET in addition to homo-FRET leading to the reduced fluorescence anisotropy. The complex formation between DOX and DNA origami as well as the interactions of the DOX excited states at different [DOX]/[bpDNA] ratios are summarized in Figure 8 to visualize the results of the study.

**Figure 8.**
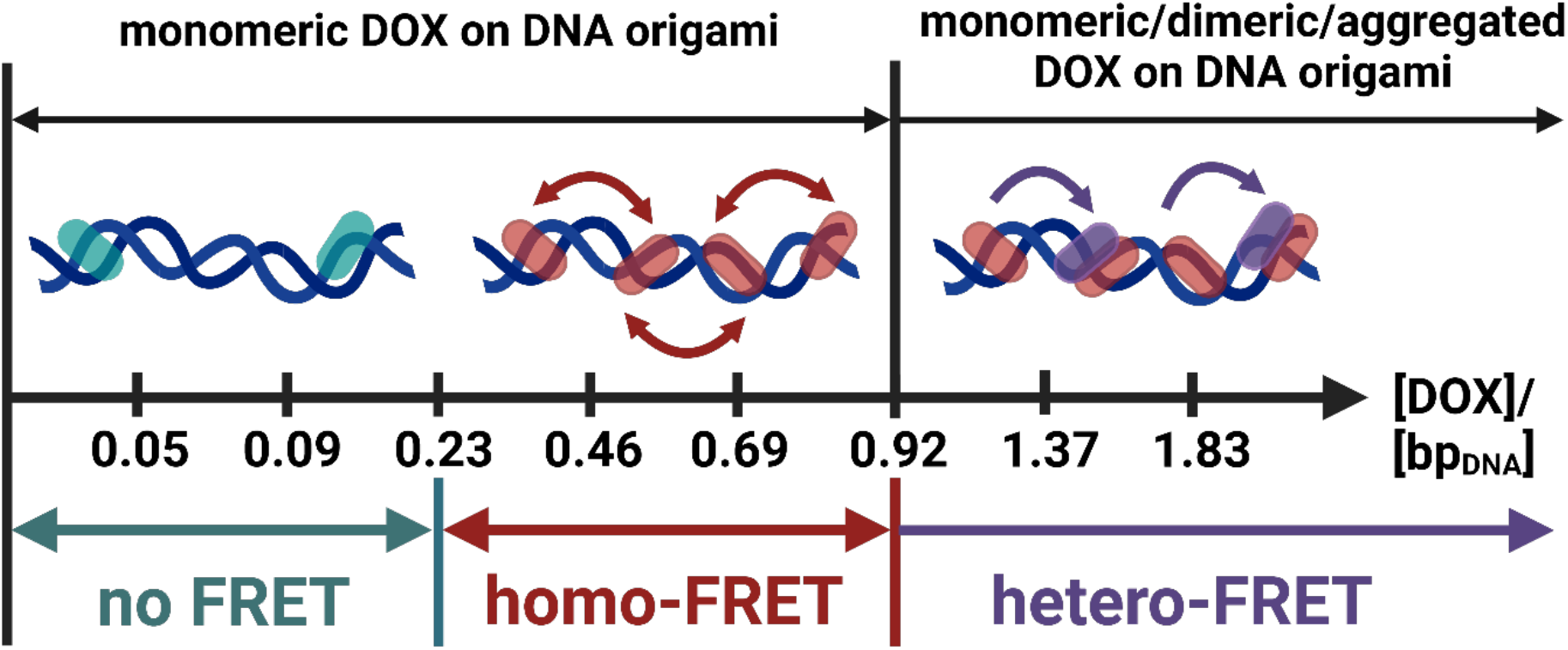
Schematic representation of the DOX binding modes in purified samples and the observed energy transfer mechanisms at different [DOX]/[bpDNA] ratios. DOX is monomeric and complexed with DNA at [DOX]/[bpDNA] < 0.92, while DOX dimers/aggregates start to appear at [DOX]/[bpDNA] > 0.92. Intercalated DOX molecules are far from each other in the DNA origami matrix at [DOX]/[bpDNA] < 0.23 resulting in no FRET. At 0.23 < [DOX]/[bpDNA] < 0.92, DOX molecules are packed closer to each other, thus facilitating the homo-FRET between identical DOX monomers. Once the intercalation sites are occupied at [DOX]/[bpDNA] > 0.92, the excess DOX forms dimer/aggregates on top of the already bound DOX molecules. The energy can be transferred from the monomeric DOX to the dimers at a close distance, thus allowing for the hetero-FRET process.

## 3. Conclusion

Applying non-destructive fluorescence anisotropy techniques to verify preparation conditions for DOX-DON drug carrier systems ensures their quality and provides relevant and comprehensive information about the DOX and DON interaction to better understand their behavior and properties for any in further uses. Fluorescence anisotropy confirmed DOX-DNA complex formation upon DOX-loading and measurements of polarized time-resolved fluorescence complemented the results of the steady-state ones, thus, revealing the hindered rotation of the DOX in DONs. The fluorescence anisotropy of free DOX and DON-bound DOX differs drastically, allowing us to assess the purity of DOX-DONs by detecting the presence of free DOX, and determining the threshold for loading ratios that would require purification for the practical applications. This may allow for further implementation in the quality control of DOX-DON formulations and tracking of drug release out of such nanocarriers. Moreover, fluorescence anisotropy gives an idea about the density of DOX packing through the homo-FRET detection in DONs or other carriers. DOX aggregation in the DON matrix was revealed for the highest loading ratios tested, and the applied purification did not remove those aggregates completely. This observation is rather important and useful for the further development of such drug delivery systems, and we hope to see these studies extended to other drug-polymer binding systems. We believe that the techniques presented here will also have applications well beyond the current DOX-DON studies.

## 4. Methods

### Doxorubicin-Loading of DNA Origami

60-helix bundle (60HB) was chosen as the DON to study the DOX-DON interaction.^[80]^ Details of 60HB production and characterization are presented in SI. Doxorubicin hydrochloride (DOX, 579.99 g mol^-1^, CRS, European Pharmacopoeia Reference Standard), dissolved in deionized water (10 mM stock), was stored at -20 °C. To reduce aggregation before the buffer exchange, 60HB in 1× folding buffer (1x FOB comprising of 1× Tris-Acetate-EDTA buffer (1× TAE buffer, containing 40 mM Tris, 20 mM acetic acid, 1 mM EDTA), 20 mM MgCl_2_, 5 mM NaCl) underwent overnight incubation (30 °C, 600 rpm), before exchanging 1× FOB to deionized water, following an adapted protocol by Kielar et al. (SI).^[10]^ DON concentration was estimated via absorbance (SI). DOX loading reactions in water contained 2 nM DONs and DOX concentrations (0.5 – 20 µM) spanning eight different [DOX]/[bp_DNA_] loading ratios (Table 1).

After vortexing, the loading reactions were incubated for an hour minimum at room temperature in the dark. The free DOX was removed via spin-filtration (adapted from Ijäs et al.^[31]^): The loading reaction (480 µL) was spun in a pre-rinsed Amicon Ultra 0.5 mL Centrifugal Filter with 100 kDa MWCO (Merck Millipore, 6 000 g, 6 min). After two washes with water (480 µL, 6 min, 6 000 g), the purified DOX-DONs were recovered from inverted filter units (1 000 g, 2 min) and diluted with water (460 µL) to their approximate original volume before storage at 4 °C. DOX-DONs’ structural integrity were checked via agarose gel electrophoresis and transmission electron microscopy (**Figure SI1, 2**). Absolute DOX concentration in the purified DOX-DONs samples was determined via calibration curve for absorption at 543 nm using a Varian Cary 50 UV-Vis Spectrophotometer (quartz cuvette with 1 cm pathlength, **Figure SI3, Table SI4**).

### Steady-State Spectroscopy

Using an UV–Vis spectrophotometer Shimadzu UV-3600 (Kyoto, Japan), the absorption spectrum of free DOX for calculation of spectral overlap was recorded in the range of 350-700 nm. The fluorescence spectra were obtained using a FLS-1000 spectrofluorometer from 500 to 800 nm (Edinburgh Instruments, U.K.). The excitation wavelength was 483 nm and the excitation spectra were monitored at 600 nm.

The steady-state anisotropy was measured by the same device using two polarizers, one located between the excitation source and the sample and the second between the sample and the emission detector. The steady-state anisotropy (*r*) spectrum was calculated automatically by the software of the spectrofluorometer based on four measurements *I*_VV_(*λ*), *I*_VH_(*λ*), *I*_HH_(*λ*), and *I*_HV_(*λ*), where subscript V refers to the vertical and H to the horizontal polarization for excitation and emission in this order. Measurements were taken within the 500-800 nm range at *λ*_exc_ = 483 nm for most of the samples, and at *λ*_exc_ = 500, 520, 540, 560 and 570 nm for samples with homo-FRET. Fluorescence anisotropy was averaged from the range 590-650 nm where the fluorescence intensity was high enough and the r value was constant (**Figure SI5**).

The distance at which the energy efficiency of FRET is 50%, *R*_0_, is calculated in Å by **Equation 1** ^[57]^

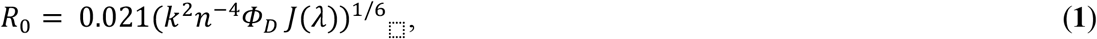

where Φ_D_ is the fluorescence quantum yield of the donor in the absence of the acceptor, *n* is the refractive index of the environment where the FRET takes place, *k*^*2*^ is the orientation factor, and *j(λ)* is the overlap integral of the absorption spectrum of the acceptor and the emission spectrum of the donor. It is given by **Equation 2**

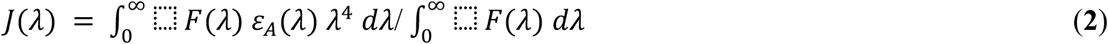

where F(*λ*) is fluorescence intensity of the donor and *ε*_*A*_ is the extinction coefficient of the acceptor. The spectral overlap *j(λ)* between the absorption and emission spectra was calculated using Origin software. DOX peak extinction coefficient at 494 nm, *ε*_494_ = 13 008 M^−1^ cm^−1^, was obtained from the absorption spectrum, in good agreement with 13 500 M^-1^ cm^-1^.^[81]^

The steady-state anisotropy *r* was calculated from **Equation 3**

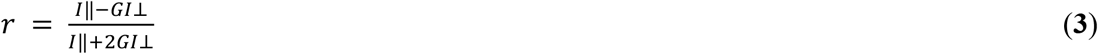

where *I‖* is the fluorescence intensity parallel to the polarization of the excitation, *I ⊥* - fluorescence intensity perpendicular to the polarization of the excitation and *G* is a correction factor to account for different transmission and detection efficiencies for parallel and perpendicular polarization. The steady-state anisotropy depends on the initial (limiting) anisotropy *r*_0_, the fluorescence lifetime *τ* and the rotational correlation time *θ* of the fluorophore according to **Equation 4** (Perrin equation)^[57]^

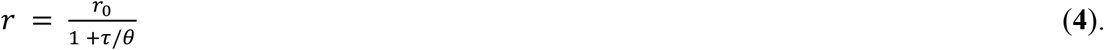

### Time-Resolved Spectroscopy

Fluorescence decay curves were measured using a time-correlated single photon counting (TCSPC) system (PicoQuant, GmBH) containing a PicoHarp 300 controller and a PDL 800-B driver. A pulsed laser LDH-P-C-485 excited the sample at 483 nm with an optical pulse width of ∼100 ps. The signals were detected with a microchannel plate photomultiplier tube (Hamamatsu R2809U). The system was also equipped with a film polarizer between the excitation source and the sample and a polarizing Glan–Taylor prism between the sample and the emission detector. Fluorescence decays were monitored at the fluorescence maximum of DOX, i.e., 600 nm. The instrumental response function (IRF) was measured separately at 483 nm and used for deconvolution analysis of the fluorescence decays followed by their fitting by sum of exponentials (**Figure SI8, Equation 5**)

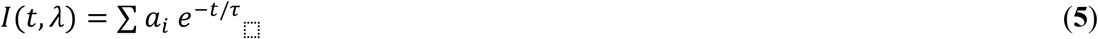

where τ is the fluorescence lifetime, and *a*_*i*_ is the amplitude (pre-exponential factor). The amplitude-averaged lifetimes are calculated from the parameters obtained via fitting by **Equation 6**^[82]^

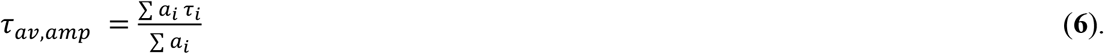

Polarization-resolved fluorescence decays were obtained when fluorescence emission passed through a polarizing beam splitter cube (Glan-Taylor prism) which could be rotated to separate the two orthogonal polarization components (parallel and perpendicular) before reaching the detector. Experimental time-resolved fluorescence anisotropy decays were produced using the parallel and perpendicular fluorescence decays, *I‖* and *I* ⊥, combined as follows in **Equation 7**

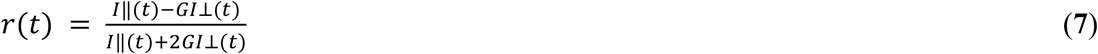

where G corresponds to the detector sensitivity ratio towards vertically and horizontally polarized light.

The fluorescence anisotropy decays were fitted with a hindered rotation single exponential decay model according to **Equation 8**

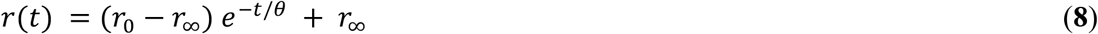

where *r*_*∞*_ accounts for hindered rotation^[57]^ for DOX complexed with DONs and is zero in case of free rotation for free DOX in water.

Molecular volume and diameter of DOX was calculated using the Stokes-Einstein relation assuming the molecule to be spherical in **Equation 9**

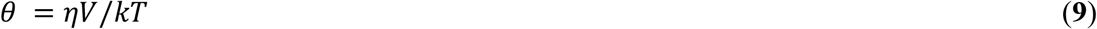

where *η* is viscosity of the media, *k* is the Boltzmann constant, and *T* – the temperature of the solution.

## Supporting Information

The article is accompanied by the Supporting Information file.

## Acknowledgments

We acknowledge financial support by the Emil Aaltonen Foundation, the Sigrid Jusélius Foundation, the Jane and Aatos Erkko Foundation, ERA Chair MATTER from the European Union’s Horizon 2020 research and innovation programme under grant agreement No 856705, European Research Council Consolidator Grant (PADRE, 101001016), Academy of Finland (project no. 323669, flagship GeneCellNano, project no. 320165, PREIN flagship programme), and the Finnish Cultural Foundation. K.S. would like to acknowledge BBSRC funding BB/R004803/1. We acknowledge the provision of facilities and technical support by Aalto University at OtaNano - Nanomicroscopy Center (Aalto-NMC). A.K. would like to thank the Biohybrid Materials Group under Prof. Mauri Kostiainen at Aalto University for their support and access to their facilities. Table of contents and Figure 1 were created with BioRender.com.

## Notes

### Competing Interest Statement

The authors have declared no competing interest.

