## Supporting Information for "Fluorescence Anisotropy Analysis of the Interaction between Doxorubicin and DNA Origami Nanostructures"

**Supporting Information: Table of Content**

**Material and Methods: DNA origami sample preparation and characterization; buffer exchange to DI water, chemical structure of DOX**

The **design and folding protocol** for the 60-helix bundle (60HB) DNA origami was used as reported by Linko et al.[1] The 60HB was folded in a 50 µL one-pot reaction containing the p7249 scaffold strand (Tilibit) at 20 nM final concentration and staple strands in 10× excess (Integrated DNA technologies) in a 1× folding buffer (1× FOB comprising of 1× Tris-Acetate-EDTA buffer (1× TAE buffer, containing 40 mM Tris, 20 mM acetic acid, 1 mM EDTA), 20 mM $MgCl_2$, 5 mM NaCl). The folding mixture was annealed with a thermal annealing ramp using an Applied Biosystems ProFlex PCR system by Thermo Fisher Scientific (cooling from 65 °C to 59 °C with a decreasing rate of 1 °C per 15 min, and from 59 °C to 40 °C decreasing 0.25 °C per 45 min).

Excess staple strands were removed via polyethylene glycol **(PEG) precipitation**, as previously reported by Stahl et al.[2] The folded DNA origami solution (~20 nM) was diluted with 1× FOB to ~5 nM and mixed with PEG precipitation buffer (15% (w/v) PEG 8000, 1× TAE, 505 mM NaCl) in a 1:1 ratio. After centrifugation at 14 000 g for 30 min, the supernatant was carefully removed and the DNA origami pellet was resuspended in the desired volume of 1× FOB overnight at 30 °C at 600 rpm using an Eppendorf ThermoMixer C, before storing it at 4 °C.

The **DNA origami concentration c** was approximated from the absorbance A at 260 nm using a BioTek Eon Microplate Spectrophotometer (2 µL sample volume, Take3™ micro-volume plate) and taking into account the corresponding buffer blank via the Lambert-Beer law ($A = \varepsilon c l$, here: pathlength $l = 0.05$ cm).[3,4] The estimated molar extinction coefficient ε at 260 nm was $0.91 \times 10^8$ $M^{-1}$ $cm^{-1}$ based on the number of hybridized and non-hybridized nucleotides of the 60HB.[5]

To verify the removal of free staple strands and the successful folding, **agarose gel electrophoresis** was performed. A 2% agarose gel was cast in 1× TAE with 11 mM $MgCl_2$ and ethidium bromide (final concentration: 0.46 µg $mL^{-1}$). The sample was diluted with 6× gel loading dye solution (Sigma Aldrich) before loading onto the gel. The gel was run at 90 V for 50 min in 1× TAE, 11 mM $MgCl_2$ as Running buffer and DNA bands were visualized under UV light using a BioRad ChemiDoc MP imaging system (SI1).

Visual verification was obtained by imaging via **Transmission Electron Microscopy** (TEM) using a FEI Tecnai 12 Bio-Twin electron microscope (120 kV acceleration voltage). The sample (3-5 µL) was deposited on plasma cleaned (20 s, Fischione Instrument NanoClean Model 1070) Formvar carbon-coated copper grids (FCF-400-CU, Electron Microscopy Sciences), similarly as described by Castro et al.[6] After 3-4 min of incubation, the sample was blotted away and negatively stained with 2% (w/v) uranyl formate solution (pH-adjusted with 25 mM NaOH) by first immersing the grid in a 5 µL stain droplet and immediately blotting it away, then in a 20 µL droplet and letting it incubate for 45 s. After the final blotting, samples were dried for at least 30 min before imaging or storage (SI2).

For the DNA origami **buffer exchange** from 1x FOB to deionized water before DOX-loading, the protocol from Kielar *et al.* was adapted[7]: In short, after pre-rinsing the Amicon Ultra 0.5 mL Centrifugal Filter with 100 kDa molecular weight cut-off (MWCO; Merck Millipore) with water (12 000 g, 5 min), DONs in 1x FOB (175 µL) were centrifuged (6 000 g, 10 min). After washing with water (314 µL, 6 000 g, 10 min), concentrated DONs in water were recovered by centrifuging the inverted filter unit (1 000 g, 2 min). Diluted with water (110 µL), the DONs concentration was determined via absorbance as described before.
The DOX-loading reactions were prepared as described in the main text, using 10 mM DOX stocks (**Figure SI0**).

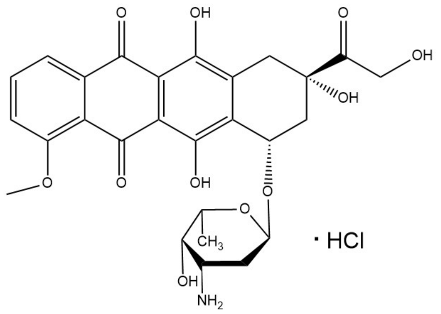

**Figure SI0.** Chemical structure of Doxorubicin hydrochloride.

**SI1: Agarose gel electrophoresis of DONs and DOX-DONs**

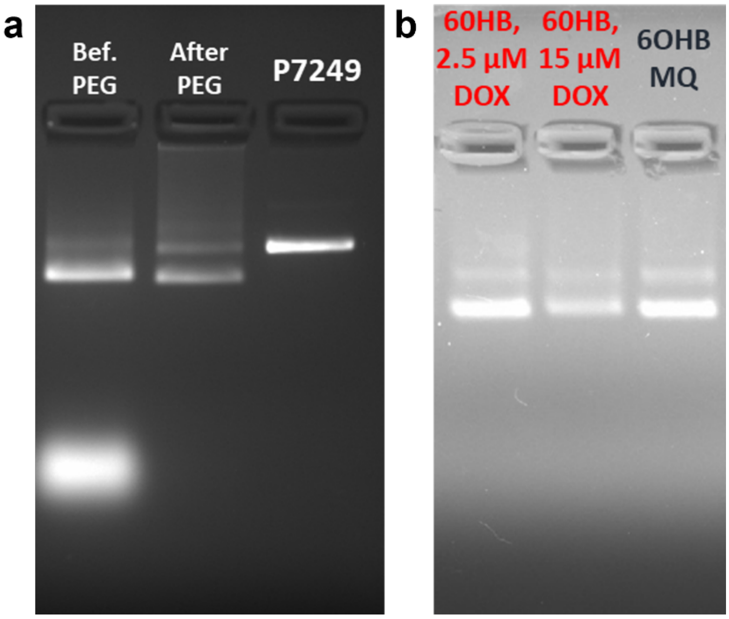

**Figure SI1**. (a) 2% Agarose gel with 0.46 µg $mL^{-1}$ ethidium bromide of freshly folded 60HB before PEG-purification to remove free excess staples (lane 1), after PEG-purification (lane 2) and the scaffold used for folding (7249 nt, lane 3). (b) 2% Agarose gel with 0.46 µg $mL^{-1}$ ethidium bromide of purified 60HB nanostructure loaded with 2.5 µM and 15 µM DOX and a reference of 60HB in deionized water (For both gels: 1x TAE, 11 mM $MgCl_2$ as Running buffer, 90V, 50 min).

The PEG-purification had successfully removed excess staple strands from 60HB folding. Folded 60HB migrated further on the agarose gel than the scaffold used. DOX-loading into the DNA nanoparticles did not influence its stability or running speed.

**SI2: Transmission Electron Microscopy images of DONs and DOX-DONs**

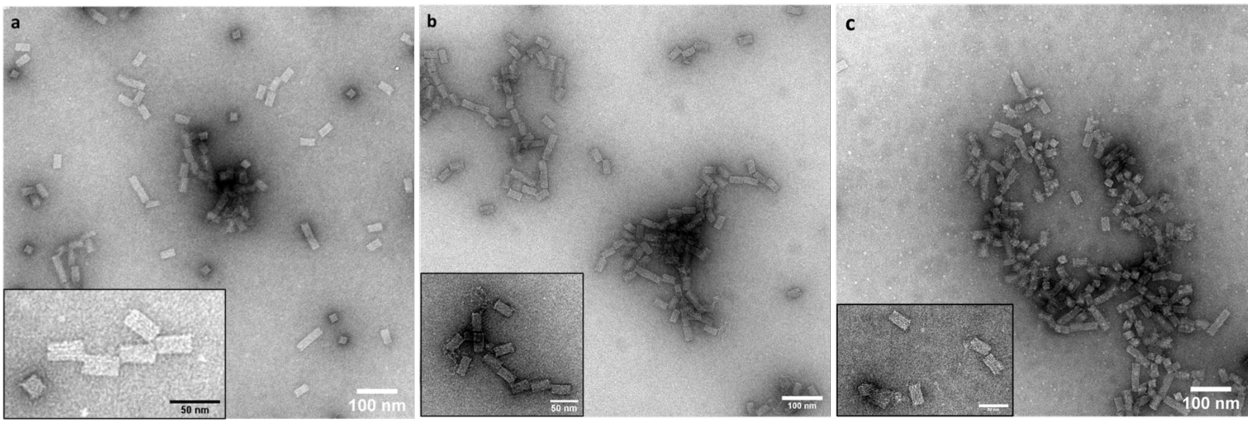

**Figure SI2**. Transmission Electron Microscopy images (120 kV, negatively stained with 2% uranyl formate solution) of (a) 60HB in FOB, (b) 60HB in deionized water, and (c) 60HB loaded with 15 µM DOX after purification (as inset: zoomed in structures with a 50 nm scale bar).

DNA origami nanostructures in FOB looked all intact. Depending on which of its surfaces the 60HB landed on during fixation, they appeared either square or oblong. Some stacking of the 60HB lead to formation of elongated oblong structures. For 60HB in water, some unraveling of a few structures was visible as some free yarn-like clumps, likely due to spin-filtration. Otherwise, they appeared stable and intact. DOX-loaded 60HB looked stable and were not deformed.

### SI3: Calibration curve of absorbance vs. DOX concentration in water

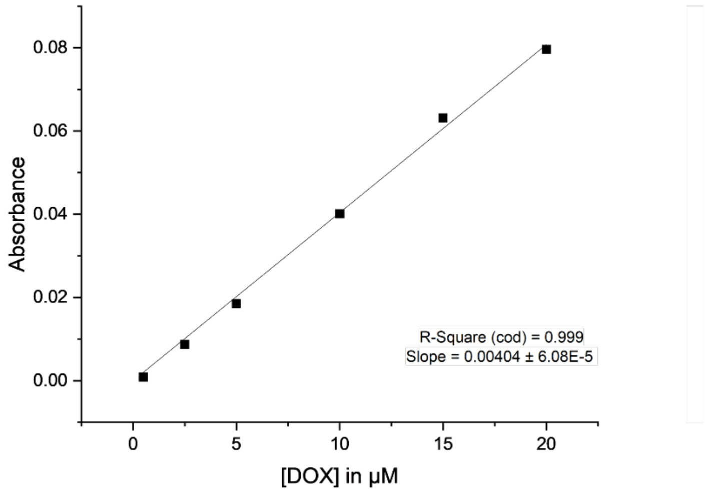

**Figure SI3**. Calibration curve for DOX concentrations in µM in deionized water at an absorbance of 543 nm (Isosbestic point).

The calibration curve (intercept set to 0) was used to determine the amount of DOX left in the purified DOX-DONs samples (**Table SI4**).

**SI4: Concentration of DOX after purification in the DOX-DONs**

**Table SI4**. Concentration of DOX [μM] in DOX-DONs before and after purification. The DOX concentration after purification was determined using the calibration curve from **Figure SI3** and considering the volume of dilution after spin-filtration.

| Loading ratio: $[DOX]/[bp_{DNA}]$ | 0.05 | 0.09 | 0.23 | 0.46 | 0.69 | 0.92 | 1.37 | 1.83 |
|---|---|---|---|---|---|---|---|---|
| **Before** purification: **DOX [μM]** | 0.5 | 1 | 2.5 | 5 | 7.5 | 10 | 15 | 20 |
| **After** purification: **DOX [μM]** | 0.35 | 0.81 | 1.42 | 2.16 | 3.04 | 3.08 | 3.94 | 6.47 |

The DOX concentration in purified DOX-DONs was reduced compared to the initial loading concentration because the spin-filtration removed free DOX/aggregates.

### SI5: Averaging of steady-state fluorescence anisotropy

The fluorescence anisotropy was measured from 495 to 800 nm (except for the excitations at longer wavelengths for Figure 6). For anisotropy values (*r*), they were averaged from the range of 590-650 nm as highlighted by the vertical dashed red lines, because the intensity of fluorescence was high enough with low noise level in that region. Moreover, the anisotropy was constant within that range, while the noise level and signal instability increased beyond the range.

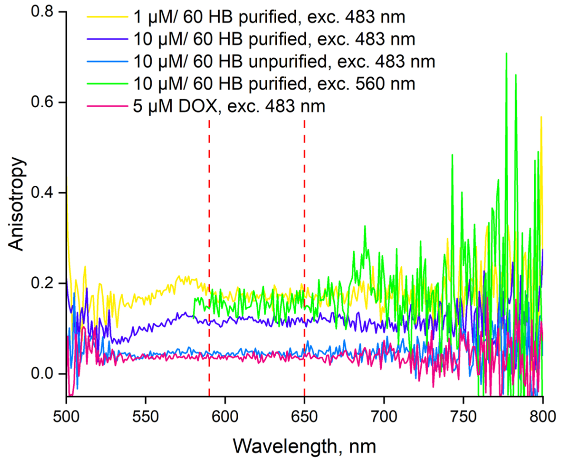

**Figure SI5**. Examples of steady-state fluorescence anisotropy spectra calculated by FLS-1000 software. Dashed red lines show the range used for averaging to obtain the final *r* values.

### SI6: Excitation and Emission spectra of DOX-DONs

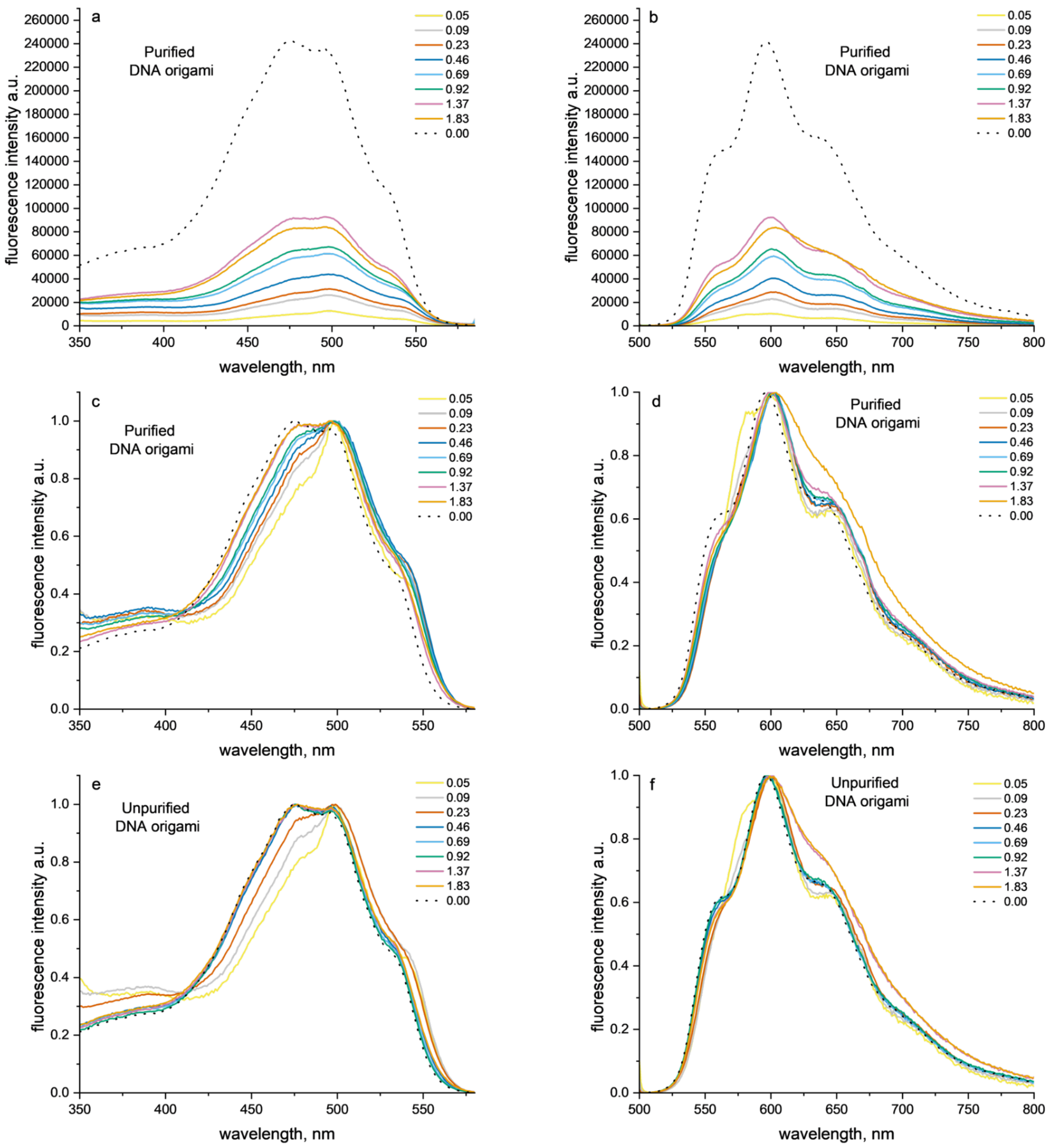

**Figure SI6**. Unnormalized and normalized excitation (a, c, e) and emission (b, d, f) spectra of free DOX in deionized water (5 µM), purified (a-d) and unpurified (e, f) DOX-DONs at different [DOX]/$[bp_{DNA}]$ loading ratios. $\lambda_{exc}$ = 483 nm, $\lambda_{det}$ = 600 nm.

### SI7: Comparison of normalized fluorescence decays of purified and unpurified DOX-DONs with free DOX

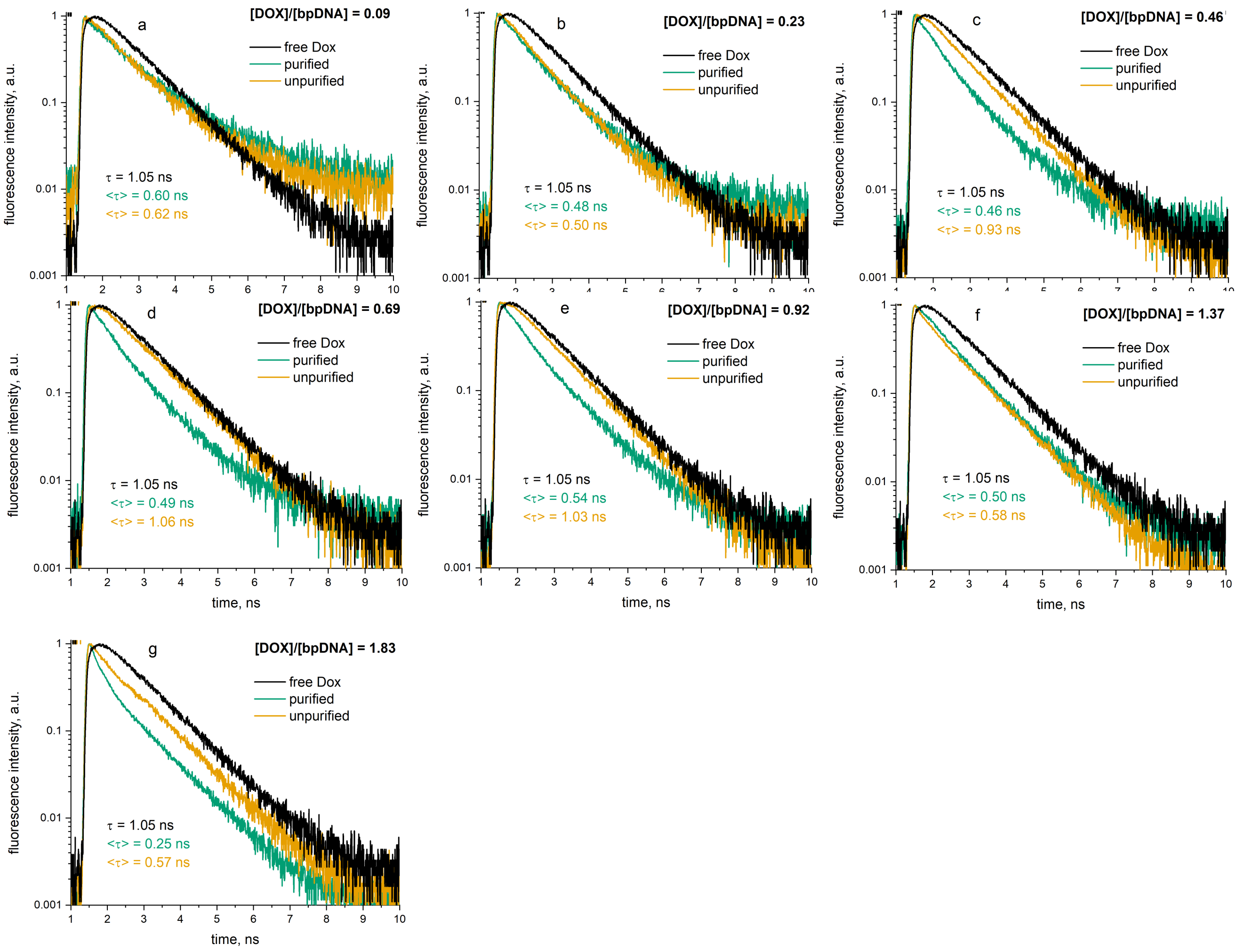

**Figure SI7**. Comparison of normalized fluorescence decays of free DOX (5μM) in deionized water with purified and unpurified DOX-DONs at different [DOX]/[$bp_{DNA}$] ratios, including their respective amplitude-weighted fluorescence lifetimes τ.

### SI8: Representative fluorescence decay fits with IRF and residuals

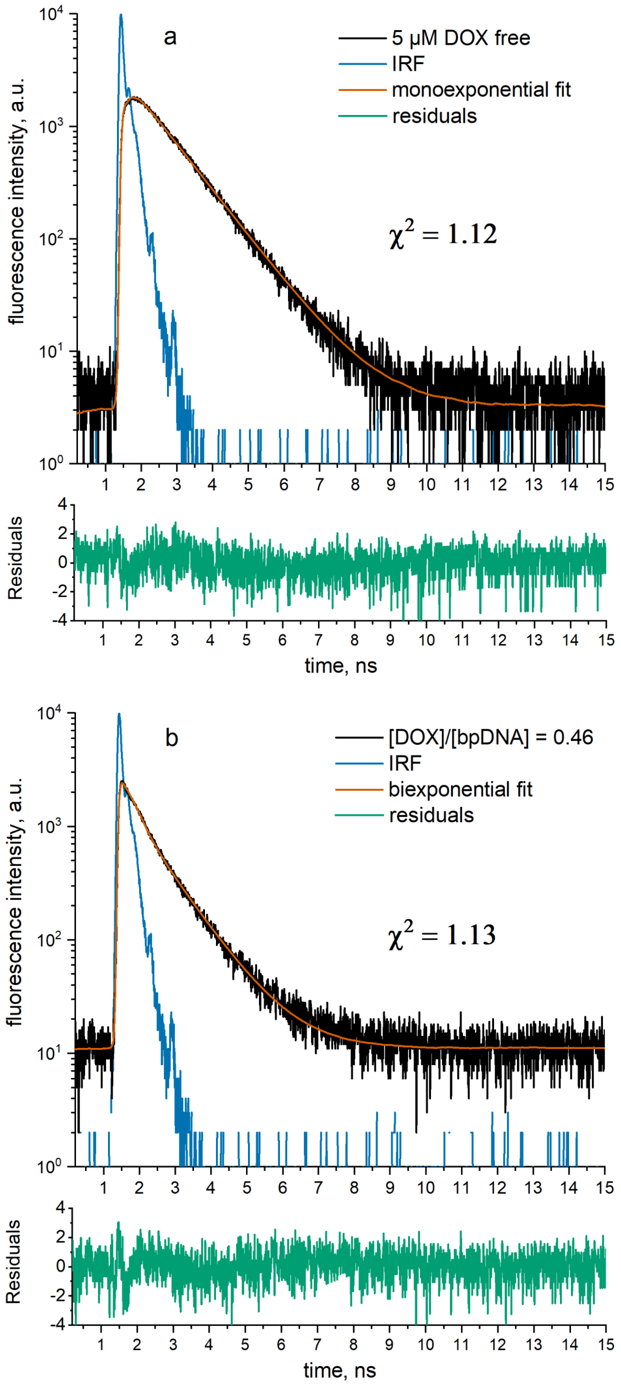

**Figure SI8**. Representative fluorescence decays of free DOX with monoexponential fit (a) and purified DOX-DON complex with biexponential fit (b) in deionized water with IRF and residuals, $\chi^2$- goodness of fit.

### SI9: Fluorescence lifetime components and contributions for DOX-DONs and free DOX

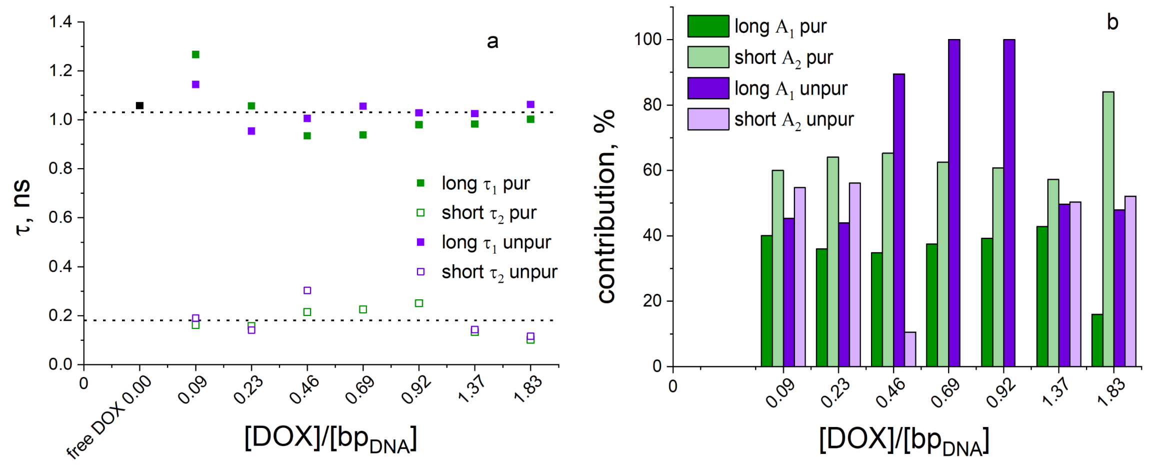

**Figure SI9**. Fluorescence lifetime components $\tau_1$ and $\tau_2$ (a) and their contributions A1 and A2 (b) obtained from the biexponential fitting of fluorescence decay curves for purified (green) and unpurified DOX-DONs (purple) at different [DOX]/[$bp_{DNA}$] loading ratios. The result of monoexponential fitting of free DOX (5 µM, black; Figure SI7) is also shown for comparison. The amplitude-averaged lifetimes are calculated from the parameters obtained from the fitting as $\tau_{av,amp} = \frac{\Sigma_i A_i \tau_i}{\Sigma_i A_i}$, where are the $A_i$ is amplitude or pre-exponential factor for each lifetime component.

### SI10: Results of the biexponential fitting for fluorescence decays

**Table SI10**. Results of biexponential fitting (Methods, Equation 2) for fluorescence decays of free DOX, purified, and unpurified DOX-DONs represented in Figure SI7.

| **Sample** | **purified** | | | | **unpurified** | | | |
|---|---|---|---|---|---|---|---|---|
| **[DOX]/ [$bp_{DNA}$]** | $\tau_1$ | **$A_1$, %** | $\tau_2$ | $\chi^2$ | $\tau_1$ | **$A_1$, %** | $\tau_2$ | $\chi^2$ |
| **0.00** Free DOX | 1.06±0.02 | 100 | na | 1.12 | - | - | - | |
| **0.09** | 1.27±0.06 | 40 | 0.16±0.05 | 1.09 | 1.14±0.06 | 45 | 0.19±0.05 | 1.05 |
| **0.23** | 1.06±0.05 | 36 | 0.16±0.03 | 1.13 | 0.95±0.02 | 44 | 0.14±0.02 | 1.09 |
| **0.46** | 0.93±0.04 | 35 | 0.21±0.02 | 1.16 | 1.00±0.01 | 89.5 | 0.30±0.11 | 1.07 |
| **0.69** | 0.94±0.04 | 37.5 | 0.23±0.03 | 1.08 | 1.06±0.01 | 100 | na | 1.10 |
| **0.92** | 0.98±0.03 | 39 | 0.25±0.02 | 1.11 | 1.03±0.01 | 100 | na | 1.13 |
| **1.37** | 0.98±0.02 | 43 | 0.13±0.03 | 1.13 | 1.02±0.01 | 50 | 0.14±0.01 | 1.09 |
| **1.83** | 1.00±0.02 | 16 | 0.10±0.01 | 1.16 | 1.06±0.02 | 48 | 0.11±0.03 | 1.13 |

### SI11: Fluorescence decay of free DOX

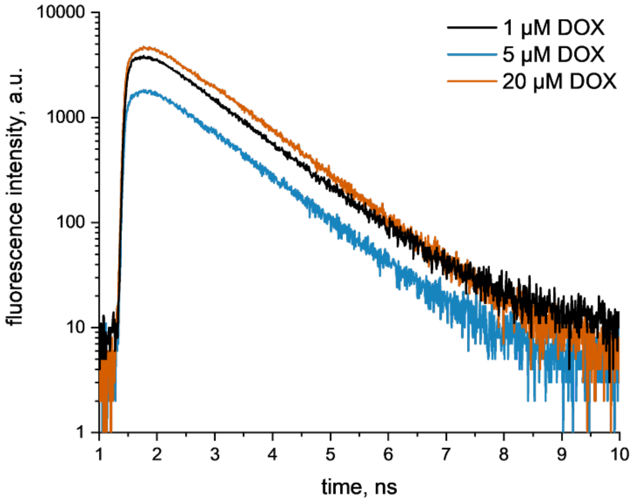

**Figure SI11**. Fluorescence decays of free DOX in deionized water at 1, 5, and 20 µM concentrations. Fluorescence lifetimes obtained from monoexponential fitting are 1.07 ± 0.03, 1.05 ± 0.02 and 1.07 ± 0.01 with the goodness of fit $\chi^2$ of 1.1-1.2 as shown in Figure SI8a.

### SI12: Calculation of Förster radius ($R_0$) for DOX homo-FRET

The overlap integral for DOX absorption and emission was calculated.

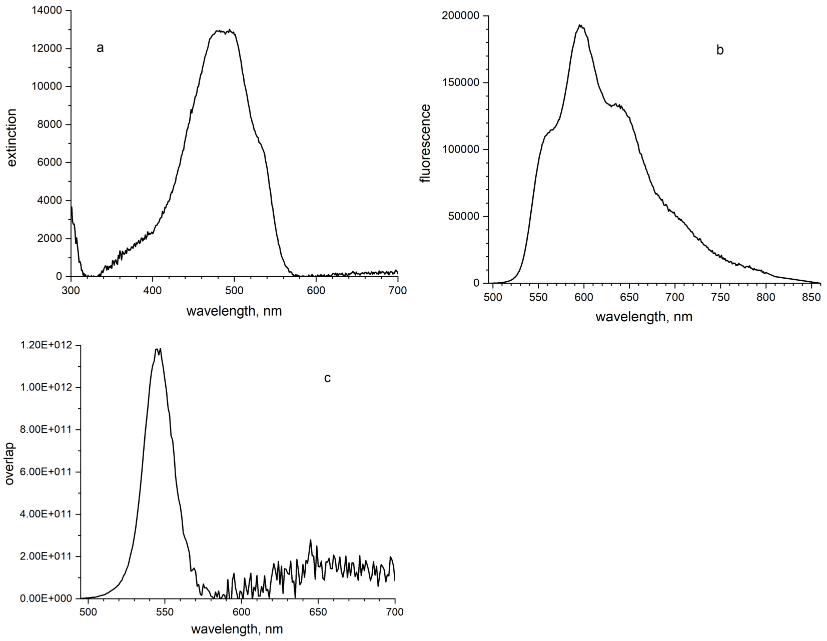

**Figure SI12**. a - Extinction spectrum $\varepsilon(\lambda)$, b - fluorescence spectrum $F(\lambda)$, and c - their overlap $J$ for 25 μM DOX in deionized water.

<u>The overlap integral</u> $J$ was calculated as

$$\int_0^\infty F(\lambda)\, \varepsilon(\lambda)\, \lambda^4\, d\lambda / \int_0^\infty F(\lambda)\, d\lambda \text{ and } d\lambda = 1 \text{ nm.}$$

The Doxorubicin extinction spectrum $\varepsilon(\lambda)$ is created from absorption by adding 0.003 and multiplying by 12,700, so the extinction peak is 13,008 l/mol/cm at 494 nm (Figure SI12a). Doxorubicin fluorescence peak is at 596 nm and integrated fluorescence $F(\lambda)$ from 495 nm to 590 nm is $2.165\times10^{11}$ (Figure SI12b). The overlap integral from 495 nm to 590 nm is $2.977\times10^{13}$ l/mol/cm nm$^4$ (Figure SI12c).

Förster radius

The Förster radius is defined as the intermolecular distance between donor and acceptor where the energy transfer efficiency is 50%. It is given by

$R_0 = 0.2108(k^2 \Phi n^{-4} J)^{1/6}$ in Å[8]

With $k^2$ = 2/3, $\Phi$ = 0.044, and $n$ = 1.33, this yields

$R_0 = 0.2108 \times (2/3 \times 0.044 \, / \, 1.33^4 \times J)^{1/6}$ Å $= 0.2108 \times (0.009375 \times J)^{1/6}$Å =
= $0.2108 \times (2.79 \times 10^{11})^{1/6}$Å = 0.2108 × 80.84 Å = 17.04 Å = 1.7 nm.

**SI13: Steady-state fluorescence anisotropy for unpurified DOX-DONs at longer excitation wavelengths**

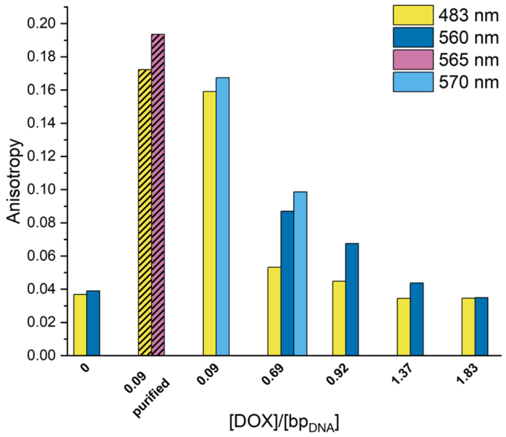

**Figure SI13**. Steady-state fluorescence anisotropy measurements at different excitation wavelengths for unpurified DOX-DONs at [DOX]/[$bp_{DNA}$] loading ratios 0.09 and 0.69-1.83, in comparison to free DOX in water (5 µM) and purified DOX-DONs of loading ratio 0.09 (shaded).

Similar as in Figure 6b (main text), shifting to longer excitation wavelengths suppresses homo-FRET which results in an increase of anisotropy values. However, compared to the purified DOX-DONs at the loading ratio 0.09, the anisotropy increase was low and did not reach its maximum. The free excess DOX in the unpurified samples dominated the anisotropy, which was unaffected by the shift to longer excitation wavelengths. Hence, the anisotropy for DOX-DONs at loading ratio 1.83 resembled that of free DOX (loading ratio 0).

### SI14: Representative parallel and perpendicular intensity decays for free DOX and purified DOX-DONs

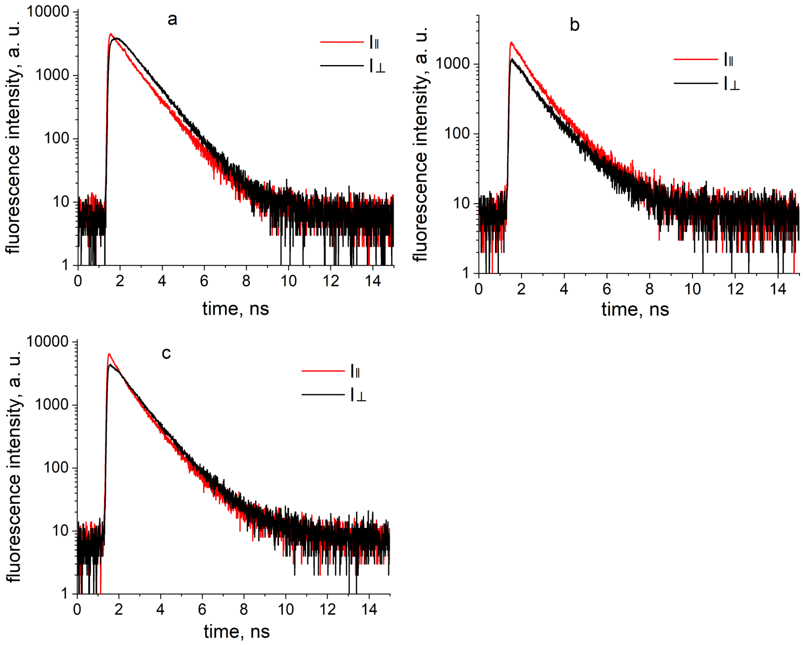

**Figure SI14**. Representative parallel and perpendicular intensity decays $I_\parallel(t)$ and $I_\perp(t)$ for free DOX in deionized water (a) and purified DOX-DONs at [DOX]/[$bp_{DNA}$] loading ratios 0.09 (b) and 0.46 (c).

**SI15: Fluorescence anisotropy decays of free DOX**

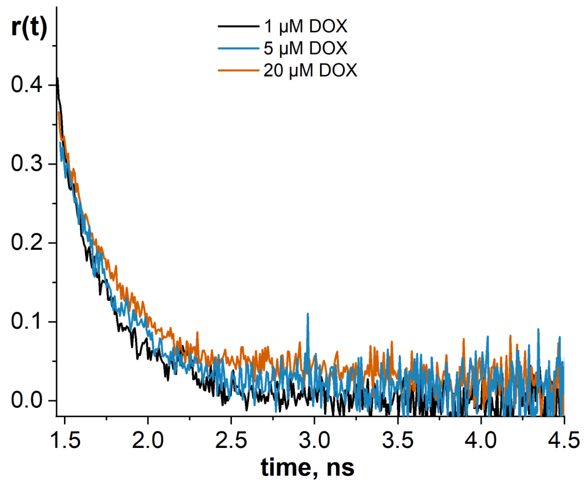

**Figure SI15**. Fluorescence anisotropy decays of free DOX in deionized water at 1, 5, and 20 μM concentration. The rotational correlation times fitted by the monoexponential model were 0.30 ± 0.01; 0.33 ± 0.01 and 0.37 ± 0.01, respectively. This yields a DOX volume of 1.21, 1.33, and 1.50 $nm^3$, and corresponding diameters of 1.32, 1.37 and 1.42 nm, assuming a spherical fluorophore. This is in good agreement with a value of 1.5 nm[9] and confirms that the signal observed originates from monomeric DOX. The difference between the rotational correlation times is below the TCSPC system resolution 0.07 ns.

**SI16: Rotational correlation times and goodness of fit for fluorescence anisotropy decays of free DOX, purified and unpurified DOX-DONs**

**Table SI16**. Rotational correlation times $\boldsymbol{\theta}$, $\boldsymbol{r_\infty}$, limiting anisotropy $\boldsymbol{r_0}$ and goodness of fit $\boldsymbol{R^2}$ from monoexponential fitting by Equation 8 in Methods for fluorescence anisotropy decays of free DOX, purified and unpurified DOX-DONs represented in Figure SI17.

| **Sample** | **purified** | | | | **unpurified** | | | |
|---|---|---|---|---|---|---|---|---|
| **[DOX]/ [bp$_{DNA}$]** | $\boldsymbol{r_0}$ | $\boldsymbol{r_\infty}$ | $\boldsymbol{\theta}$ | $\boldsymbol{R^2}$ | $\boldsymbol{r_0}$ | $\boldsymbol{r_\infty}$ | $\boldsymbol{\theta}$ | $\boldsymbol{R^2}$ |
| **0.00** Free DOX | 0.40 | 0.00 | 0.30±0.01 | 0.95 | - | - | - | - |
| **0.09** | 0.39 | 0.23 | 1.31±0.29 | 0.56 | 0.39 | 0.19 | 1.36±0.26 | 0.65 |
| **0.23** | 0.40 | 0.21 | 1.32±0.21 | 0.61 | 0.39 | 0.12 | 0.85±0.05 | 0.83 |
| **0.46** | 0.37 | 0.14 | 1.28±0.16 | 0.79 | 0.39 | 0.08 | 0.47±0.01 | 0.92 |
| **0.69** | 0.37 | 0.08 | 0.93±0.07 | 0.85 | 0.38 | 0.06 | 0.43±0.02 | 0.87 |
| **0.92** | 0.37 | 0.08 | 0.81±0.04 | 0.89 | 0.38 | 0.07 | 0.38±0.01 | 0.90 |
| **1.37** | 0.38 | 0.06 | 0.53±0.02 | 0.92 | 0.23 | 0.04 | 0.47±0.02 | 0.83 |
| **1.83** | 0.23 | 0.06 | 0.61±0.04 | 0.87 | 0.24 | 0.06 | 0.49±0.04 | 0.78 |

### SI17: Fluorescence anisotropy decays of purified and unpurified DOX-DONs

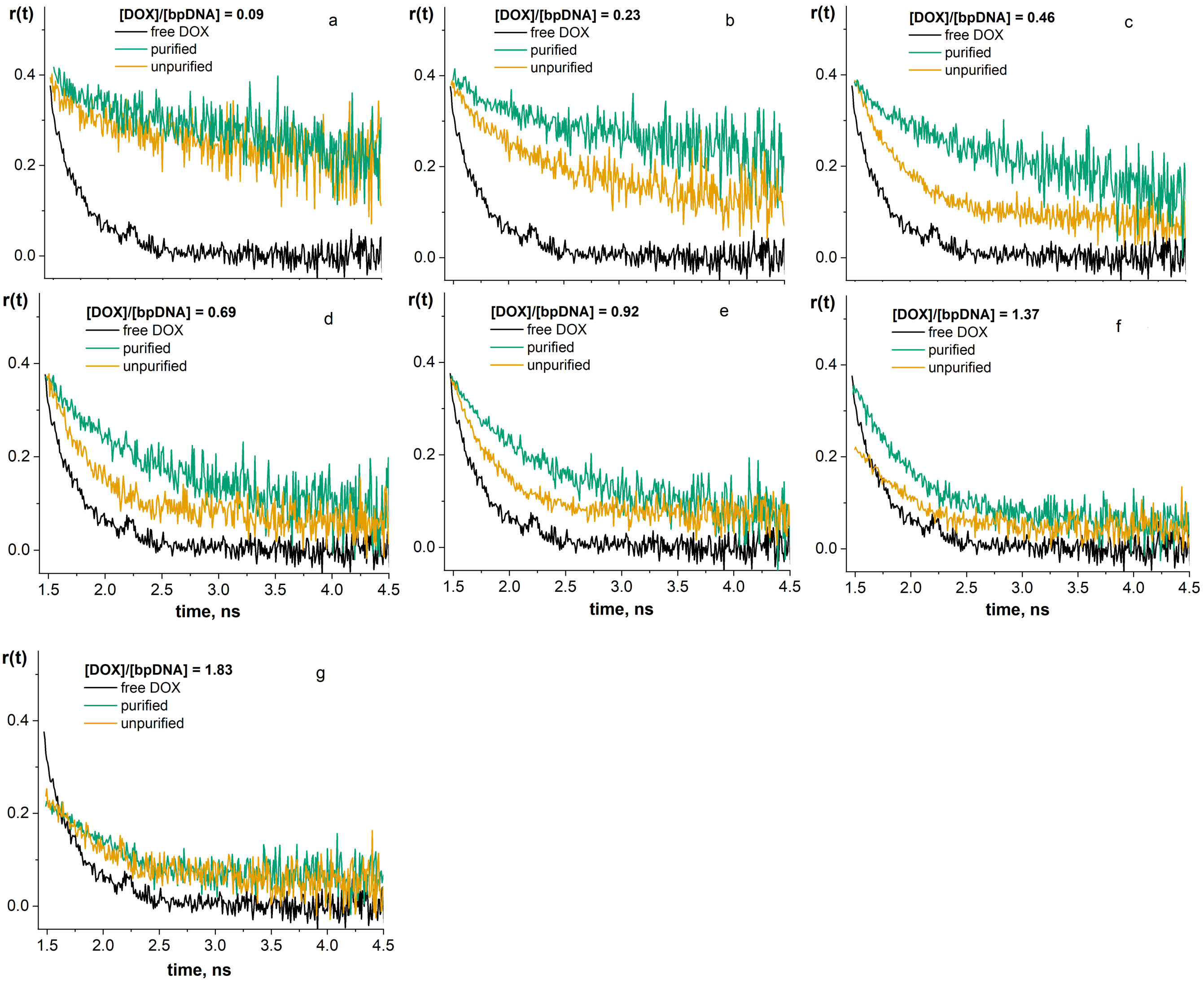

**Figure SI17.** Fluorescence anisotropy decays of purified (green) and unpurified (yellow) DOX-DONs at different [DOX]/[$bp_{DNA}$] loading ratios in comparison to that of free DOX (black). The rotational correlation times fitted by monoexponential model are presented in Table SI16.
